# Genomic features predict bacterial life history strategies in soil, as identified by metagenomic stable isotope probing

**DOI:** 10.1101/2022.09.09.507310

**Authors:** Samuel E. Barnett, Rob Egan, Brian Foster, Emiley A. Eloe-Fadrosh, Daniel H. Buckley

## Abstract

Bacteria catalyze the formation and destruction of soil organic matter, but the bacterial dynamics in soil that govern carbon (C) cycling are not well understood. Life history strategies explain the complex dynamics of bacterial populations and activities based on tradeoffs in energy allocation to growth, resource acquisition, and survival. Such tradeoffs influence the fate of soil C, but their genomic basis remains poorly characterized. We used multi-substrate metagenomic DNA stable isotope probing to link genomic features of bacteria to their C acquisition and growth dynamics. We identify several genomic features associated with patterns of bacterial C acquisition and growth, notably genomic investment in resource acquisition and regulatory flexibility. Moreover, we identify genomic tradeoffs defined by numbers of transcription factors, membrane transporters, and secreted products, which match predictions from life history theory. We further show that genomic investment in resource acquisition and regulatory flexibility can predict bacterial ecological strategies in soil.

## Introduction

Soil dwelling microorganisms are essential mediators of terrestrial C cycling^1–5^, yet their immense diversity^6,7^ and physiological complexity, as well as the mazelike heterogeneity of their habitats^8–11^, make it difficult to study their ecology *in situ*. Life history theory has been proposed as a framework for predicting bacterial activity in soils^12–15^. Life history theory^16,17^ explains the ecological properties of organisms based on their energy allocation to growth, resource acquisition, and survival^3,14,18–20^. A fundamental aspect of this framework is that life history traits impose ecological tradeoffs that constrain fitness with respect to environmental properties^16,17^. For example, tradeoffs between bacterial growth rate and yield are thought to constrain bacterial activity with respect to environmental variability^18^. Such tradeoffs can influence C fate by controlling the amount of C mineralized to CO_2_ or converted into microbial products that become SOM^18^. As a result, the accuracy of global C-cycling models can be improved by including information about microbial ecological strategies^21–23^. Unfortunately, bacterial life history traits resist *in situ* characterization, and experiments with cultured strains often ignore the complex microbe-microbe and microbe-environment interactions that occur in soil^24^.

In a previous study^15^, we quantified the dynamics of C acquisition and growth for diverse soil dwelling bacteria by performing a multi-substrate DNA stable isotope probing (DNA-SIP) experiment that tracked nine different C sources, which varied in bioavailability, through the soil food web over a period of 48 days (Fig. S1). Through this approach we demonstrated that Grime’s C-S-R life history framework explains significant variation in bacterial growth and C acquisition dynamics in soil^15^. We used these data to calculate several parameters that describe patterns of resource acquisition and growth. Resource *bioavailability* was determined as the average bioavailability of the ^13^C-labeled C sources assimilated by taxa. *Maximum log_2_ fold change* (max LFC) was determined as the maximal change in differential abundance of taxa in response to C input. *Latency* of C assimilation was determined for taxa as the difference in time between maximal ^13^C mineralization and earliest ^13^C-labelling for a given C source. Latency changes in proportion to the likelihood that taxa engage in primary assimilation of ^13^C directly from a C source, or secondary assimilation of ^13^C following microbial processing. Here we have sought to identify genomic features that underlie bacterial life history traits linked to the C-S-R framework.

Since the majority of soil dwelling bacteria remain uncultivated and poorly described^25,26^, there is great utility in identifying genomic features that predict the ecological strategies of bacteria^27^. Genomic features of life history strategies have been identified in marine bacteria^28^ and proposed for soil dwelling bacteria^29^. Genomic features associated with growth, resource acquisition, and survival are of particular interest when assessing life history tradeoffs^13,14,30,31^. Numerous genes control such quantitative traits, however, and it is difficult to predict these complex traits *de novo* from genomic data. We hypothesized that life history strategies impose tradeoffs that alter genomic investment in the gene systems (*i.e*., numbers of genes devoted to a particular system) that govern quantitative traits linked to growth, resource acquisition, and survival. We predicted that these tradeoffs would manifest in gene systems that control transcriptional regulation, membrane transport, secreted enzyme production, secondary metabolite production, motility, attachment, osmotic stress response, and dormancy. We linked genomic investment in these systems to patterns of resource acquisition and growth for soil dwelling bacteria by performing metagenomic analysis of ^13^C-labeled DNA (metagenomic-SIP) derived from our previous multi-substrate DNA-SIP experiment.

Metagenomic-SIP allowed us to link ^13^C-labeled contigs and metagenome-assembled genomes (MAGs) to patterns of resource acquisition and growth as they occurred within soil (Fig. S1). For metagenomic-SIP, we selected eight ^13^C-labeled samples from the prior experiment, because these 8 samples were enriched in genomes of taxa whose resource acquisition and growth dynamics represented extremes in the C-S-R life history framework (Fig. S1, S2). This strategy, by diminishing the confounding contribution of genomes from organisms having intermediate life-history strategies, facilitates identification of genome features that underlie life history tradeoffs. We took three approaches to analyzing these metagenomic-SIP data, each increasing in complexity: (*i*) a ^13^C-labeled contig-based approach to assess whether genome feature enrichment correlates with resource acquisition and growth parameters at community scale, (*ii*) a ^13^C-labeled MAG approach to assess whether genome feature enrichment correlates with resource acquisition and growth parameters for discrete taxa, and (*iii*) a ^13^C-labeled MAG approach to assess tradeoffs between genome features predicted from the C-S-R framework.

The third approach was designed to identify bacterial life history strategies by characterizing tradeoffs between genomic investment in regulatory flexibility and resource acquisition, as predicted from the C-S-R framework^30,31^. We chose to assess genomic investment in regulatory flexibility as the number of transcription factors (TF) relative to total gene number (TF:gene). Environmental variability will favor high TF:gene because TF regulate gene expression in response to changes in the cellular environment^32^. We chose to assess genomic investment in resource acquisition as the number of genes encoding secreted enzymes (SE), secondary metabolite biosynthetic pathways (SM), and membrane transporters (MT). SE and SM are required for acquisition and control of extracellular resources. MT are required for resource uptake and their function provides the physiological foundation for the concept of the copiotrophy-oligotrophy continuum^12,33–35^. The C-S-R framework describes tradeoffs with respect to resource acquisition and environmental variability^30,31^. Competitors (C) have high investment in resource acquisition and favor intermediate levels of environmental variability. Stress tolerators (S) have low investment in resource acquisition and are disfavored by temporal variability. Ruderals (R) have low investment in resource acquisition and are favored by high levels of temporal variability. On the basis of this framework, we predicted a tradeoff whereby investment in resource acquisition (SE + SM) would be highest relative to investment in MT for intermediate levels of regulatory flexibility (TF:gene) and lowest at both high and low levels of regulatory flexibility. By clustering MAGs based on these tradeoffs and comparing resource acquisition and growth parameters across clusters, we demonstrate the ability of these genomic features to predict bacterial life history strategies.

## Results and Discussion

### Identification of ^13^C-labeled contigs with metagenomic-SIP

We used metagenomic-SIP to enrich for DNA from ^13^C-labeled bacteria and to identify ^13^C-labeled contigs, thereby linking genomic content to C acquisition. Overall, we recovered between 5 × 10^8^ and 1.3 × 10^9^ reads in each metagenome library after quality control (Table S1). Co-assembly generated over 1.2 × 10^6^ contigs that were >1000 bp long, of which 639,258 were ^13^C-labeled in at least one treatment (>5X coverage in the ^13^C-treatment library and >1.5-fold increased coverage relative to the corresponding ^12^C-control library; Table S1). After normalizing for sequencing depth, the number of genes annotated from ^13^C-labeled contigs in each treatment was positively correlated with the number of ^13^C-labeled OTUs (Fig. S3; Pearson’s *r* = 0.795, *p*-value =0.018), as expected. The phylum representation observed for ^13^C-labeled contigs differed somewhat from that observed for ^13^C-labeled OTUs as determined by 16S rRNA sequencing (Fig. S4). This difference could be due to loss of some contigs from ^13^C-labeled metagenomic libraries on the basis of genome G + C content or due to differences in annotation methodologies used in metagenomic and 16S rRNA based methods (see Supplementary Results).

### Genomic features of ^13^C-contigs explain variation in resource acquisition and growth dynamics

We first tested whether the targeted genomic features explained variation in resource acquisition and growth dynamics at community level, as assessed across the entire collection of ^13^C-labeled contigs (Fig. S5, Fig. S6) and ^13^C-labeled OTUs observed from each ^13^C-labeled treatment (Supplementary Dataset). This contig-based approach is meaningful because ^13^C source identity had a large and significant effect on the identity of ^13^C-labeled taxa, with this variation driven by the overall dynamics of ^13^C-assimilation and growth, as previously described^15^. Three of the eight genomic features we examined explained significant variation in resource acquisition and growth dynamics (Fig. 1). Methyl-accepting chemotaxis protein genes (MCP) were positively correlated with max LFC (Pearson’s r = 0.954, p-value = 0.002; Fig. 1a), indicating that these genes are frequent in taxa that increase relative abundance dramatically in respond to new C inputs. In addition, MT (Pearson’s r = 0.907, p-value = 0.015) and osmotic stress response (OS) genes (Pearson’s r = 0.938, p-value = 0.004) were both positively correlated with C source bioavailability (Fig. 1b, c).

**Figure 1:**
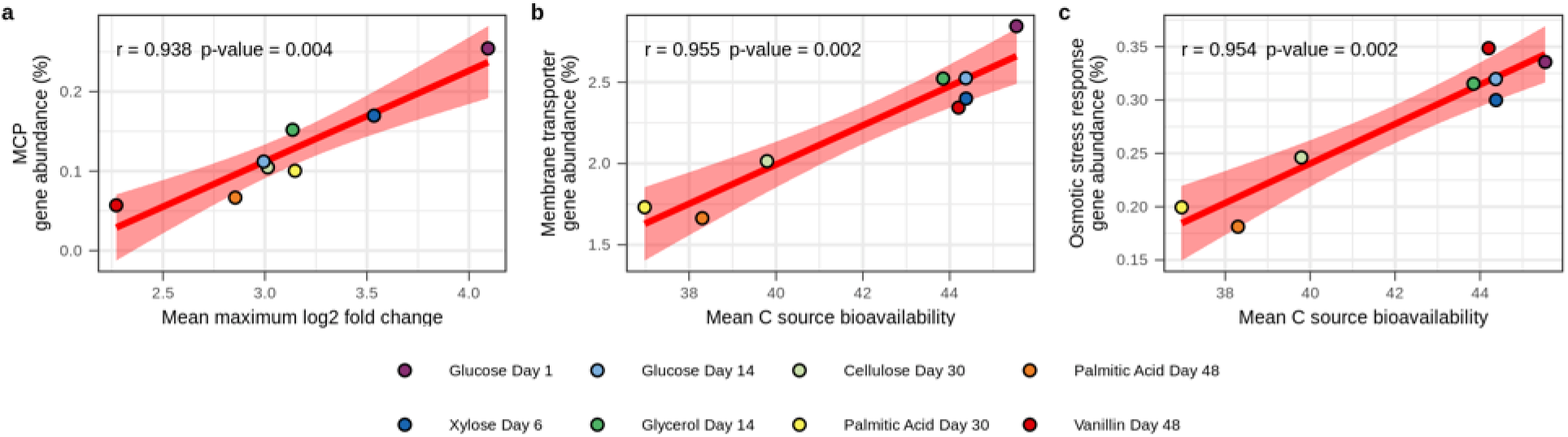
Genomic features of ^13^C-labeled contigs correlate with activity characteristics of ^13^C-labeled OTUs. **a)** Abundance of methyl-accepting chemotaxis protein (MCP) genes correlates positively with the mean maximum log2 fold change (Max LFC) of the ^13^C-labeled OTUs. **b)** Abundance of membrane transporter (MT) genes correlates positively with the mean bioavailability of C sources acquired by the ^13^C-labeled OTUs. **c)** Abundance of osmotic stress response genes (OS) correlates positively with the mean bioavailability of C sources acquired by the ^13^C-labeled OTUs. In all cases, the abundance is calculated as the percent of protein coding genes in ^13^C-labeled contigs that are annotated within the genomic feature. Red lines represent linear relationships with shading indicating the 95% confidence intervals. Pearson’s *r* and *p*-values are provided. *p*-values are adjusted for multiple comparisons using the Benjamini-Hochburg procedure (*n* = 8).

Soil consists of a complex matrix^36,37^ in which microbial access to C is limited by spatial and temporal variability^38,39^. Moisture is a major determinant of resource availability in soils, controlling soil matrix conductivity and tortuosity, and thereby regulating rates of diffusion^40–43^ as well as sorption/desorption kinetics^44^. For these reasons soil moisture is a major determinant of bacterial activity in soils^45–47^. While resource concentration is a major determinant of bacterial growth kinetics in aquatic environments, bioavailability is a major determinant of bacterial growth kinetics in soil^15^. Bioavailability, defined as the ability of a resource to cross the membrane, is determined in soil by solubility, sorption dynamics, and soil moisture^8,48,49^. High bioavailability C sources (*e.g*., glucose, xylose, and glycerol) are highly soluble, less likely to be sorbed to soil minerals, readily available for membrane transport, and their availability to cells governed primarily by diffusive transport as limited by soil moisture^35^. These substrates are degraded rapidly and so elevated concentrations are ephemeral in soils^50^. Hence, to compete effectively for highly bioavailable C sources, bacteria must exploit ephemeral periods when their resources are present in high concentration. Low bioavailability C sources (*e.g*., cellulose and palmitic acid), in contrast, cannot be transported directly across the membrane until transformed by extracellular microbial products such as secreted enzymes^3,31,51^ or biosurfactants^52^. These substrates are typically insoluble in soils and degraded over a span of weeks, months, or even years. Hence, to compete effectively for low bioavailability C sources soil dwelling bacteria must invest in resource acquisition, by manufacturing extracellular products that facilitate access to insoluble particulate materials.

Chemotactic bacteria can move through soil pore water and water films, allowing preferential access to C sources detected by MCP^53,54^. MCPs are a dominant chemoreceptor family shared by diverse bacterial phyla^55,56^, and they are widely recognized as directing chemotaxis^56,57^. Our finding that MCP genes increase in proportion to the max LFC of bacterial taxa (Fig. 1a), suggests that chemotaxis is an important determinant of fitness for bacteria whose relative abundance increases dramatically during ephemeral periods of high resource availability. Similar explosive population dynamics are expected for organisms having a ruderal strategy as described in Grime’s C-S-R framework^30^. Hence, we hypothesize that chemotaxis is adaptive in soils for growth-adapted bacteria that compete for ephemeral resources whose availability is driven by high environmental variability, and that MCP gene count is a genomic feature that can help identify soil dwelling bacteria having this life history trait.

MT activity regulates resource uptake, and transporter kinetics have been described as a key determinant of copiotrophic and oligotrophic life history strategies in aquatic environments^33–35,58^. Hence, membrane transport is likely a key determinant of bacterial life history strategies in soil. We show that high MT gene frequency correlates with the ability of soil bacteria to acquire high bioavailability C sources (Fig. 1b). We hypothesize that high MT gene count is adaptive for bacteria that compete for ephemeral, highly bioavailable C sources. In soil, high MT gene count is likely indicative of more copiotrophic bacteria with copiotrophs encompassing a wide diversity of life history strategies including both ruderals and competitors as defined by Grime’s framework^30^. We also hypothesize that low MT gene count is likely an indicator of oligotrophic bacteria that compete for less bioavailable C sources in soil, with low MT gene frequency indicating a tendency towards resource specialization.

OS genes are affiliated with several cellular systems for surviving low water activity including compatible solutes, aquaporins, and ion homeostasis^59,60^. OS systems are of vital importance for microbial survival in soils due to the high variation in water activity^61,62^. We show that OS genes are more frequent in soil dwelling bacteria that acquire C from highly bioavailable C sources (Fig. 1c). Highly bioavailable C sources are transiently abundant in water filled pore space when soils are moist^63^. Soil pores dry out rapidly as moisture becomes limiting, hence we predict that OS is adaptive for bacteria that exploit resources present in water filled pore space. In contrast, bacteria using low bioavailability C sources localize preferentially to surfaces. Water films and biofilms are favored on soil surfaces^42^, buffering the organisms localized there from rapid variation in water activity. Our results suggest that OS is adaptive for soil dwelling bacteria of more copiotrophic character (*i.e*., ruderals and competitors), those that compete for high bioavailability substrates whose availability corresponds with rapid changes in water activity.

One might naively predict that OS would be a characteristic of organisms having a stress tolerant life history strategy. The observation that OS does not predict a ‘stress tolerant’ bacterial lifestyle requires us to carefully consider how we define ‘stress’ in bacterial ecology. Grime’s original framework, from plant ecology, describes plant stress as limitation for light, nutrients, and/or water, which are resources required for plant growth^30^. This plant-centric definition of stress, based on resource limitation, conflicts with the microbiological definition, in which ‘stress’ is usually interpreted as abiotic stress (*e.g*., tolerance to pH, salinity, temperature, O_2_). Those bacteria that are adapted for resource limitation are typically defined as oligotrophs. Hence, Grime’s ‘stress tolerator’ strategy, as interpreted in the proper ecological context, is indicative of bacteria having oligotrophic characteristics^15^, and not those adapted for extremes of abiotic stress (*e.g*., extremophiles). These contrasting definitions of stress are a potential source of confusion when life history theory developed for plants is applied to bacteria. We propose that a better understanding of bacterial life history theory would be provided by interpreting the ‘S’ in C-S-R as a ‘scarcity-adapted’ rather than ‘stress-adapted’.

### Genomic features of ^13^C-MAGs explain variation in resource acquisition and growth dynamics

A limitation of the contig-based analysis described above is that statistical power is low since we have only 8 treatments. Hence, we also used MAGs to evaluate associations between genomic features and activity characteristics. We recovered 27 ‘medium quality’ MAGs^64^ from the ^13^C-labeled contigs (> 50% completeness and < 10% contamination; Supplemental Dataset; Supplemental Results). We linked these MAGs to corresponding ^13^C-labeled OTUs present in the exact same ^13^C-labeled DNA sample on the basis of taxonomic annotations (assigned by GTDBtk^65^, Supplemental Dataset). For example, the ^13^C-labeled MAG Glucose_Day01_bin.1 was classified to the family *Burkholderiacea* and therefore linked to all *Burkholderiacea* OTUs ^13^C-labeled in the glucose day 1 treatment. Three MAGs did not match any OTU (Cellulose_Day30_bin.7, PalmiticAcid_Day48_bin.4, and Vanillin_Day48_bin.1), while the others matched 1–56 OTUs each. For each ^13^C-labeled MAG, activity characteristics were averaged across the matching ^13^C-labeled OTUs (Fig. S7, Supplemental Dataset). We then evaluated the number of genes associated with each genomic feature, normalized for MAG size (Fig. S8, Supplemental Dataset). As before, MT genes were positively correlated with C source bioavailability (Pearson’s r = 0.550, p-value = 0.043; Fig. 2A), and we found that TF genes (Pearson’s r = 0.881, p-value < 0.001) and secondary metabolite biosynthetic gene cluster (SMBC) abundance (Pearson’s r = 0.712, p-value = 0.001) were also positively correlated with C source bioavailability (Fig. 2b, c).

**Figure 2:**
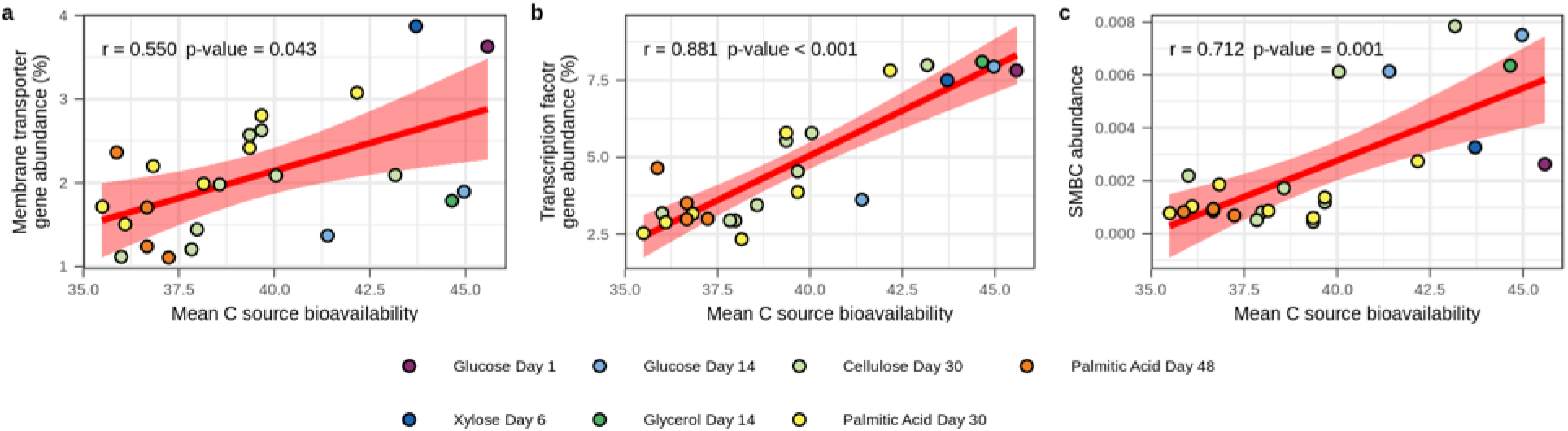
Genomic features of ^13^C-labeled MAGs correlate with activity characteristics of ^13^C-labeled OTUs taxonomically and isotopically mapped to MAGs. **a)** MT frequency, **b)** TF frequency, and **c)** SMBC abundance all correlate positively with the mean bioavailability of C sources acquired. For MT and TF, frequency is calculated as the percent of protein coding genes in MAGs that are annotated within the genomic feature. For SMBCs, abundance is the number of SMBCs divided by the number of protein coding genes in MAGs. Red lines represent linear relationships with shading indicating the 95% confidence intervals. Pearson’s *r* and *p*-values are provided. *p*-values are adjusted for multiple comparisons using the Benjamini-Hochburg procedure (*n* = 8).

Having high numbers of TF is thought to be an adaptive trait for microbes living in highly variable environments^32,66,67^. Certain taxa are known to be enriched in TF families but the evolutionary basis of variation in TF gene frequency is not well established^68^. Our finding that TF frequency correlates with C source bioavailability (Fig. 2b) suggests that growth on ephemeral C sources favors high TF, because this adaptive trait allows bacteria to respond effectively to high environmental variability. The metabolic and physiological changes induced by these TF may include previously discussed features such as MCP, MT, or OS systems. Our results support the idea that genomic investment in TF is an adaptive trait that varies with environmental variability of the ecological niche.

Secondary metabolites include a wide range of small molecules produced by organisms. Bacteria often use these molecules to interact with their environments. Examples include antibiotics that kill or prevent the growth of other organisms, signaling molecules that mediate intercellular interactions, siderophores, chelators, and biosurfactants used to access insoluble nutrients^69^. Secondary metabolites can facilitate competition for limited resources^70,71^ and they can even mediate microbial predation^72^. Production of secondary metabolites requires multiple genes often found in clusters (*i.e*., SMBCs)^73,74^. We show that SMBC frequency correlates with C source bioavailability (Fig. 2c). This finding, runs counter to the idea that secondary metabolites are important for competition on low bioavailability resources^69,75,76^. Given that this observation matches patterns observed for TF and MT we expect that SMBC are favored by conditions of environmental variability and/or resource acquisition.

### Genomic feature correlation in publicly available soil genomes and metagenomes

We observed through metagenomic-SIP that C source bioavailability correlates with MT, OS, TF and SMBC frequencies and we hypothesize that these gene frequencies are predictive of an organisms position on the copiotroph-oligotroph continuum. From this hypothesis, we predict that these genomic features should correlate in independent genomic and metagenomic datasets. We assessed these relationships in several datasets generated from a range of different soils (see Supplementary Results). Since MT were significantly associated with C source bioavailability at both community level (^13^C-labeled contigs) and genome level (^13^C-labeled MAGs), we compared the gene frequencies for MT with those of TF, OS, and SMBCs in each independent dataset. Support for a relationship between MT and both TF and OS was supported in 4 of 7 independent datasets (Fig. 3a-e). We found no correlation between MT and SMBC frequencies within any of the datasets (Fig. 3).

**Figure 3:**
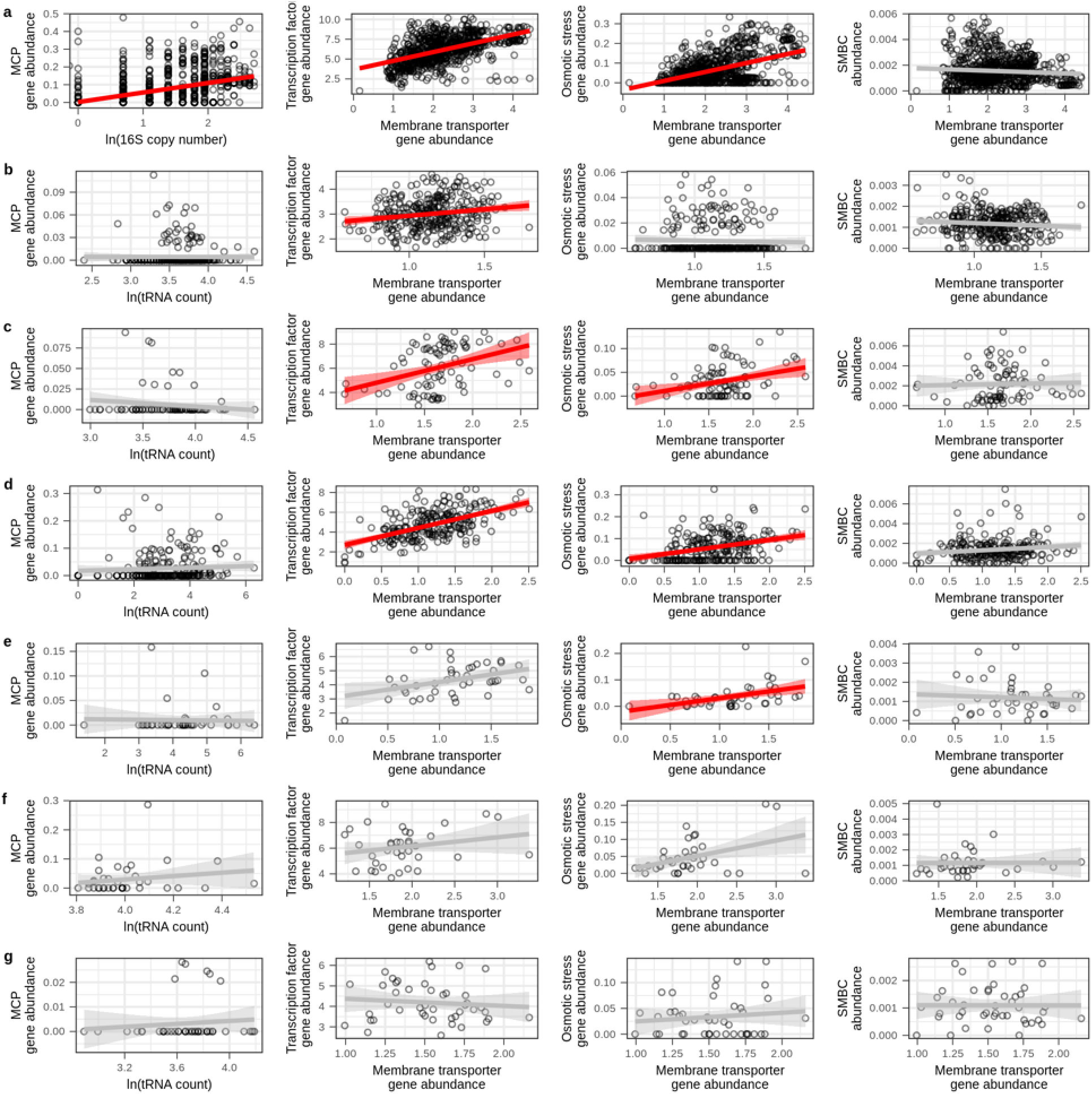
MT correlates with TF and OS in 4 of 7 independent metagenomic datasets examined and MCP correlates with log *rrn* copy number in the RefSoil database. The tRNA gene count was used as a proxy for *rrn* copy number as described in text. The datasets are **a)** RefSoil genomes, **b)** Diamond *et al*. 2019 MAGs recovered from drought simulated meadow soils, **c)** Yu *et al*. 2020 MAGs recovered from heavy DNA extracted from agricultural soils supplied with ^13^C-labeled ryegrass, **d)** Wilhelm *et al*. 2019 MAGs recovered from heavy DNA extracted from forest soils treated with either ^13^C-labeled cellulose or lignin, **e)** Wilhelm *et al*. 2021 phylobins recovered from heavy DNA fractions extracted from agricultural soil supplied with ^13^C-labeled cellulose, **f)** Zhalnina *et al*. 2018 genomes isolated from *Avena barbata* rhizosphere, and **g)** Li *et al*. 2019 MAGs recovered from rhizospheres of *Zea mays, Triticum aestivum*, and *Arabidopsis thaliana*. Red or grey lines represent the linear relationships with shading indicating the 95% confidence intervals. Red relationships are statistically significant (adjusted *p*-value < 0.05) with *p*-value adjusted for multiple comparisons within dataset using the Benjamini-Hochberg procedure (n = 4). Correlation statistics are in Supplementary Dataset.

We also observed that MCP gene counts (Fig. 1a) and predicted rRNA gene (*rrn*) copy number^15^ both correlate with max LFC when new C is added to soil. We hypothesize that these traits are linked to ruderal strategies (a subset of copiotrophs), hence we predict that *rrn* copy number should correlate with MCP gene frequency in independent datasets. We compared MCP gene frequency to the natural log of either *rrn* copy number (for RefSoil), or tRNA gene count (for reference metagenome MAGs). While the RefSoil database contains complete genomes with accurate *rrn* copy numbers, MAGs from metagenomic datasets do not provide accurate *rrn* annotations, therefore we used tRNA gene abundances as a proxy since tRNA gene count correlates with *rrn* copy number^77^. In further support of this proxy, we observed that *rrn* copy number and tRNA gene count are strongly correlated in RefSoil bacterial genomes (Pearson’s r = 0.792, p-value < 0.001; Fig. S9). The natural log of *rrn* copy number was positively correlated to MCP gene abundance across the RefSoil dataset (Fig. 3a), yet the natural log of the tRNA gene counts were not correlated with MCP gene abundance in any of the other datasets (Fig. 3b-g).

A correlational approach, as applied above, has two notable limitations. First, many of the genes in metagenomic datasets are poorly annotated. Inaccurate annotation can produce inaccurate gene counts for all of the gene systems we assessed. Second, adaptive tradeoffs between gene systems will not produce straightforward correlations, because the concept of a tradeoff implies an interaction whereby the adaptive benefit varies depending on the life history strategy of the organism^78^.

### Tradeoffs in genomic investment define life history strategies

Tradeoffs occur when the benefit of a trait in a given environment differs between two groups. For example, increases in environmental variability might tend to favor more investment in resource acquisition for oligotrophic organisms (because higher variability tends to produce higher average nutrient levels when resources are low), but less investment in resource acquisition in copiotrophic organisms (because investing in extracellular products that enable resource acquisition provides little benefit in a highly disturbed environment). To detect, among our ^13^C-labeled MAGs, tradeoffs between regulatory flexibility, resource acquisition, and membrane transport, we examined relationships between TF:gene and [SE + SM]:MT. The ratio TF:gene measures genomic investment in regulatory flexibility. The ratio [SE + SM]:MT captures genomic investment in resource acquisition relative to uptake. SM represents the sum of all genes found in SMBCs, reflecting genomic investment in secondary metabolite biosynthesis. We summed SM and SE because these features represent genomic investment in extracellular products. Groups of genomes adapted to similar life history strategies should exhibit comparable genomic investment in these gene systems. We used *k*-means clustering based on genomic investment in these gene systems to group the MAGs into three clusters that we hypothesized would represent the C-S-R strategies. We then determined whether the properties of the genomes in each cluster matched predictions from the C-S-R framework.

We observed evidence for tradeoffs in both regulatory flexibility and resource acquisition among these three clusters. TF tended to increase with total gene count (as expected), but TF:gene differed between the three clusters (Fig. 4a). When genome size was small, the three clusters differed little in TF, but as total gene count increased the clusters diverged with one cluster having less regulatory flexibility than the other two (Fig. 4a). We also observe that [SE + SM] gene counts tend to increase in proportion to MT counts in two clusters (as expected), but the other cluster, which has the highest MT counts, maintains low [SE + SM] counts (Fig. 4b). When these relationships are plotted together, we observe that one cluster tends to increase relative investment in resource acquisition ([SE + SM]:MT) along with regulatory flexibility (TF:gene), while the other two have the opposite response (Fig. 4c).

**Figure 4:**
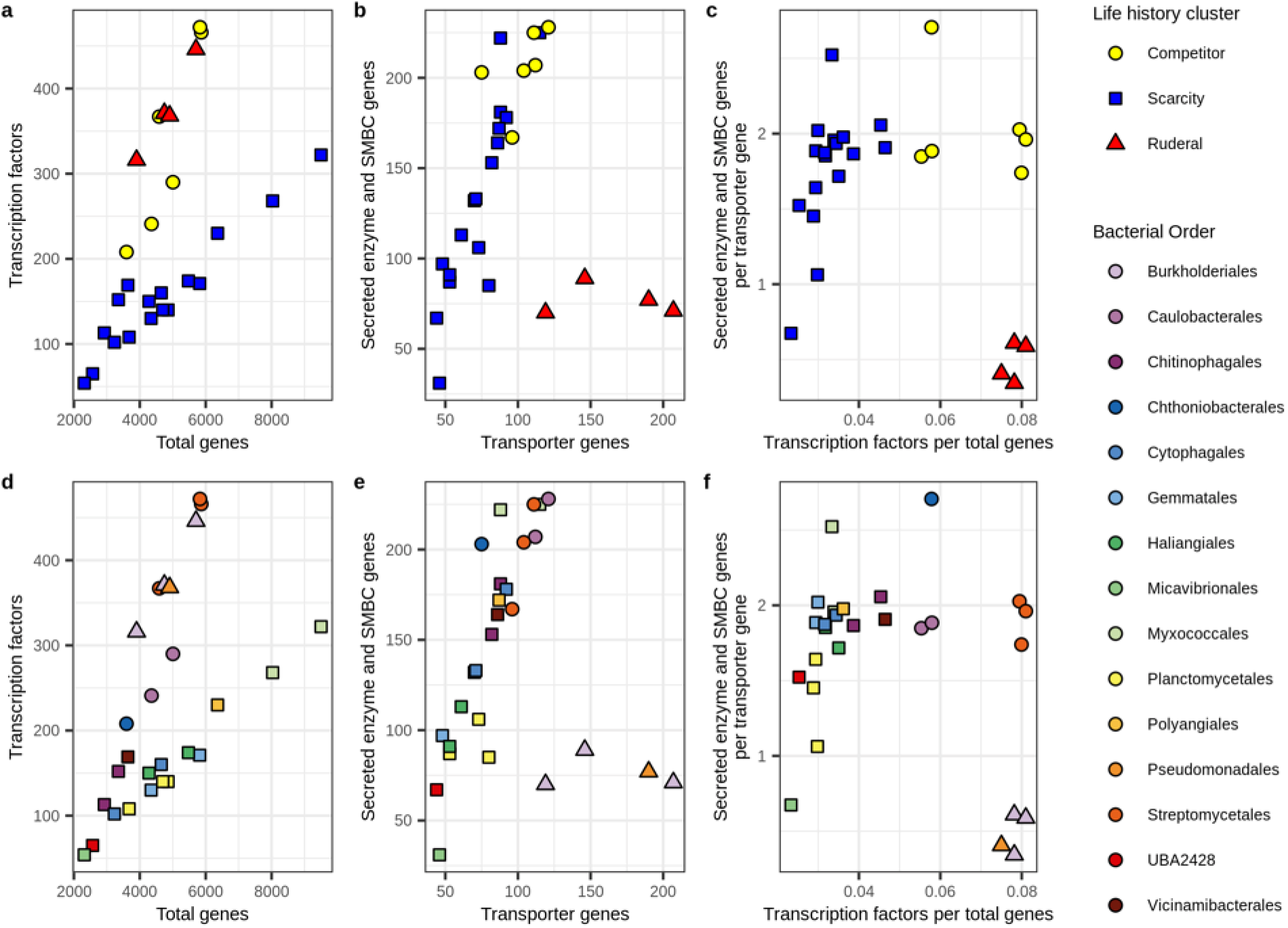
Genomic investment in gene systems can be used to cluster MAGs into life history strategies. MAGs were grouped using k-means clustering on scaled values of TF:genes and [SE + SM]:MT. **a)** Relationship between TF and total gene count. **b)** Relationship between summed SE and SM gene counts and MT, where SM indicates total genes within SMBCs. **c)** The relationship between genomic investment in resource acquisition ([SE + SM]:MT) and regulatory flexibility (TF:genes). Clusters are colored by predicted life history strategies within the C-S-R framework. **d-f)** The taxonomic identifies of the MAGs (at the order level) corresponding to panels a-c.

These three clusters demonstrate adaptive tradeoffs consistent with Grime’s C-S-R framework. The scarcity strategists (*i.e*., oligotrophs; S) have low regulatory flexibility (Fig. 4a), and generally low genomic investment in transport (Fig. 4b), but their genomic investment in resource acquisition tends to increase in proportion to regulatory flexibility (Fig. 4c). That is, scarcity strategists whose ecological niches are the most constant require little genomic investment in regulatory flexibility and resource acquisition, while those whose niches are more variable require more investment in regulatory flexibility and resource acquisition. In contrast, ruderals (R) have high regulatory flexibility (Fig. 4a), and high investment in transport (Fig. 4b), but they have low genomic investment in resource acquisition (Fig. 4b, c). Finally, the competitive strategists (C) have intermediate to high levels of regulatory flexibility (Fig. 4a), intermediate investment in membrane transport (Fig. 4b), but high genomic investment in resource acquisition (Fig. 4a) with little relationship between resource acquisition and regulatory flexibility (Fig. 4c). We expect many intermediate strategies among the C-S-R vertices, and as expected we see that scarcity specialists adapted for high levels of regulatory flexibility are difficult to distinguish from competitive specialists adapted for lower levels of regulatory flexibility.

MAGs assigned to the three clusters differ in their resource acquisition and growth dynamics consistent with the expectations of life history theory. Ruderals and competitors acquired C sources that had significantly higher bioavailability than scarcity specialists (Fig. 5a), and they also consumed a higher diversity of C sources than the scarcity specialists, and this difference was significant (Fig. 5d). Ruderals, however, had significantly higher max LFC relative to competitors indicating the ability to increase population size dramatically in response to C input (Fig. 5b).

**Figure 5:**
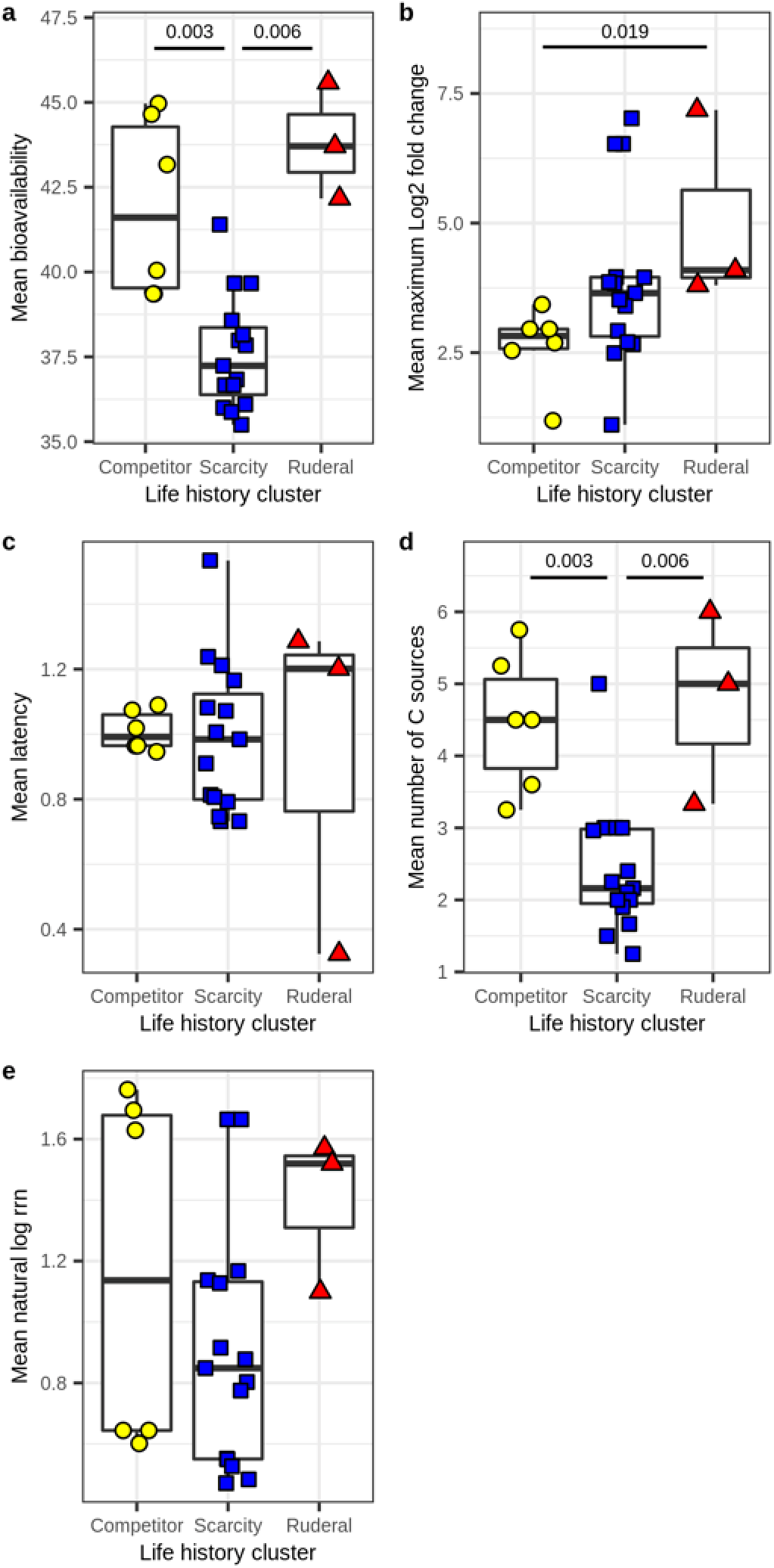
Resource acquisition and growth dynamics differ across life history strategies indicative of tradeoffs predicted from Grime’s C-S-R framework. Clusters corresponding to life history strategies were determined from *k*-means clustering based on TF:genes and [SE + SM]:MT, as previously indicated (from Fig. 4). Significance was determined by Kruskal-Wallis tests with post hoc comparisons performed using Dunn tests. **a)** Bioavailability of ^13^C sources acquired was lower for scarcity adapted MAGs than for competitor or ruderal MAGs. **b)** Max LFC was higher for ruderal MAGs than competitor MAGs. **c)** No difference was observed in latency across the three clusters. **d)** Number of ^13^C sources acquired was lower for scarcity adapted MAGs than for competitor or ruderal MAGs. **e)** No difference was observed in the natural log of *rrn* copy number across the clusters.

In terms of genomic features, we see that both ruderals and competitors have higher TF and OS gene frequencies than scarcity specialists (Fig. 6a), while only the ruderals have higher MT relative to scarcity specialists, and these differences are significant (Fig. 6a). Ruderals are distinguished from both competitors and scarcity specialists by their low investment in SE and high investment in MCP (Fig. 6a). Competitors are distinguished from both scarcity and ruderal specialists by their higher investment in adhesion (Fig. 6a). The general theme is that both ruderals and competitors have copiotrophic characteristics, but ruderals appear to be opportunists with adaptations that maximize their ability to exploit ephemeral resources, while competitors have greater genomic investment in resource acquisition. Scarcity specialists appear less well adapted for regulatory flexibility and more likely to specialize in their C sources (Fig. 5d).

**Figure 6:**
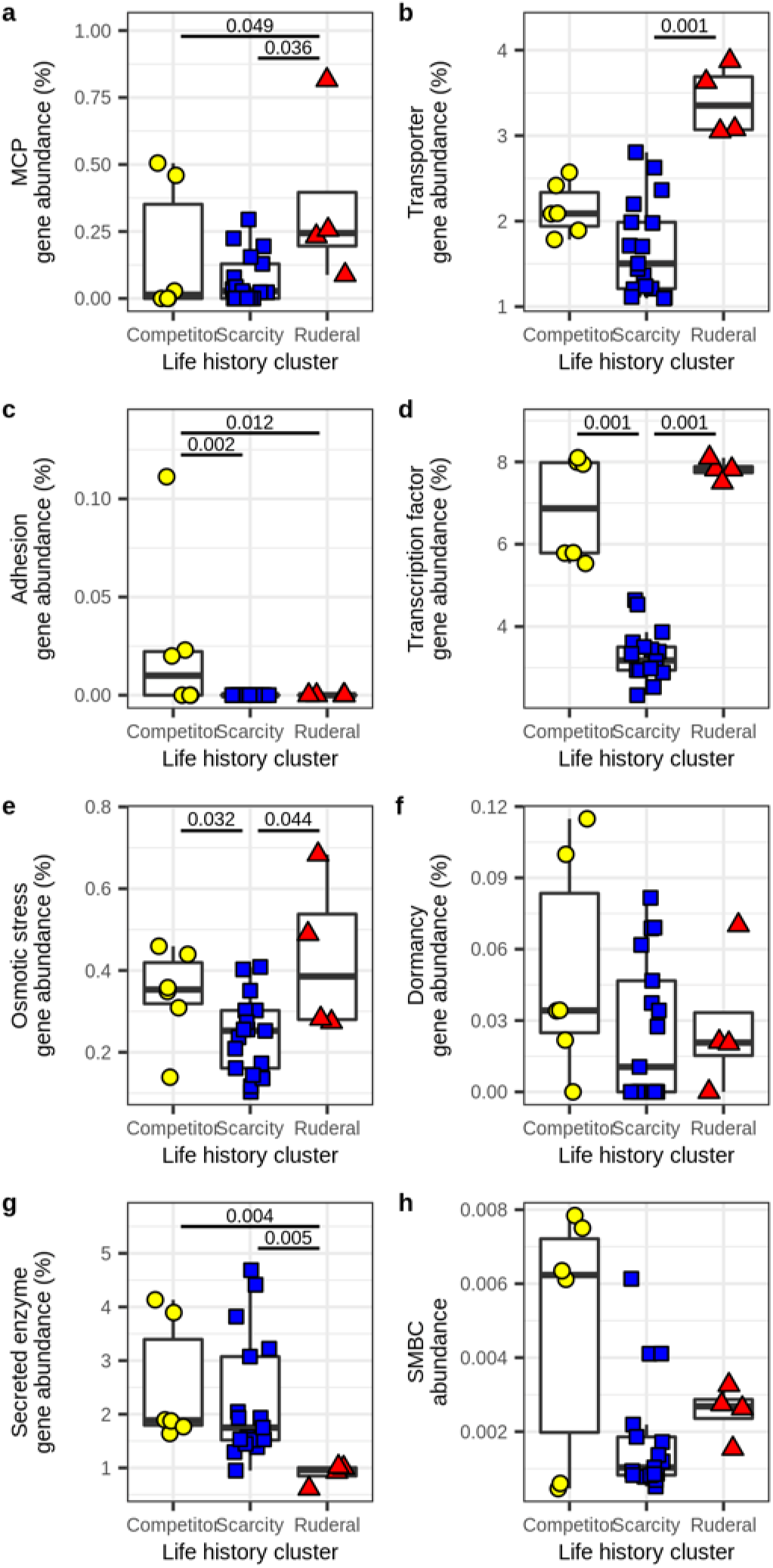
Genomic investment in gene systems differs across the three life history strategies indicative of tradeoffs predicted from Grime’s C-S-R framework. Clusters corresponding to life history strategies were determined from *k*-means clustering based on TF:genes and [SE + SM]:MT, as previously indicated (from Fig. 4). Significance was determined by Kruskal-Wallis tests with post hoc comparisons performed using Dunn tests. **a)** Ruderal MAGs have higher investment in MCP than competitor or scarcity adapted MAGs. **b)** Ruderal MAGs have higher investment in MT than scarcity adapted MAGs. **c)** Competitor MAGs have higher investment in adhesion genes than ruderal or scarcity adapted MAGs. **d)** Scarcity adapted MAGs have a lower investment in TF than ruderal or competitor MAGs. **e)** Scarcity adapted MAGs have a lower investment in OS than ruderal or competitor MAGs. **f)** There is no statistically significant difference in investment in dormancy genes across clusters. **g)** Ruderal MAGs have a lower investment in SE than competitor or scarcity adapted MAGs. **h)** There is no statistically significant difference in investment in SMBCs across clusters.

### Predicting ecological strategies from genome features

We used parameters of TF:gene and [SE + SM]:MT, defined from the three ^13^C-labeled MAG clusters described above, to predict life history strategies for RefSoil genomes. The resulting RefSoil genome clusters, predicted from these genome parameters, exhibited genomic characteristics representative of the expected life history tradeoffs (Fig. 7a-c). The relationship between TF:gene and [SE + SM]:MT is roughly triangular, as we would expect for the C-S-R framework (Fig. 7c). It is apparent that a vast diversity of intermediate life history strategies exist (Fig. 7c), and this is also an expected result since relatively few taxa will maximize adaptive tradeoffs while most will optimize adaptive traits to suit their particular ecological niche. Genomes having ruderal characteristics are enriched in the *Gammaproteobacteria* and *Firmicutes* (Fig. 7f, Fig. S10), as we would expect, though members of these phyla can be found in all three clusters (Fig. S10) owing to the vast diversity of these groups. In addition, genomes having competitive characteristics are highly enriched in the *Actinobacteria* and *Betaproteobacteria*, while genomes characteristic of scarcity specialists are enriched in the *Alphaproteobacteria* and other diverse phyla (*e.g., Verrucomicrobia, Acidobacteria, Gemmatamonadetes, Chloroflexi, etc.*) whose members are difficult to cultivate in laboratory media (Fig. 7f, Fig. S10). Most bacterial phyla are metabolically and ecologically diverse and we would not expect homogeneity among species within a phylum. In addition, previous observations show that C assimilation dynamics in soil are not well predicted by phylum level classification^15^. However, certain strategies are more common in some phyla than others, and these patterns, along with the taxonomic makeup of our MAG clusters (Fig. 5d-f) match general expectations. Furthermore, the three clusters we defined for RefSoil genomes possess patterns of genomic investment that match predictions derived from the C-S-R framework and are consistent with predictions based on the ^13^C-labeled MAGs (Fig. S11, Table S2).

**Figure 7:**
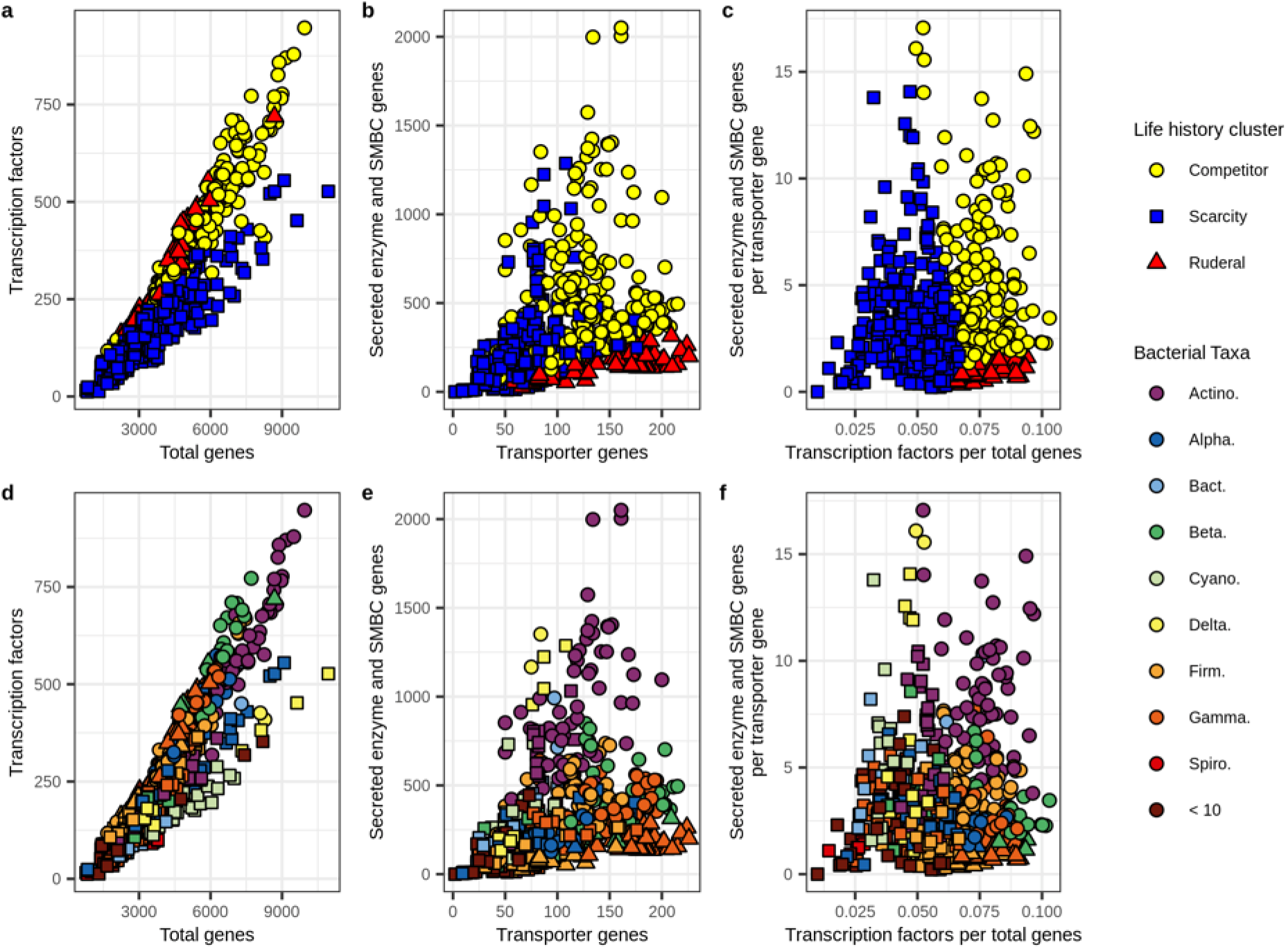
Tradeoffs in genomic features can be used to predict life history strategies from reference genomes. RefSoil bacterial genomes were clustered based on genomic tradeoffs between resource acquisition ([SE + SM]:MT) and regulatory flexibility (TF:genes) using *k*-means clustering trained on the three clusters defined for ^13^C-labeled MAGs (from Fig. 4) **a)** Relationship between TF and total gene count. **b)** Relationship between summed SE and SM gene counts and MT, where SM genes are total genes within SMBCs. **c)** The relationship between genomic investment in resource acquisition ([SE + SM]:MT) and regulatory flexibility (TF:genes). Clusters are colored by predicted life history strategies within the C-S-R framework. **d-f)** Taxonomic identifies of genomes corresponding with panels a-c (at the phylum or class level: Actino. = *Actinobacteria*, Alpha. = *Alphaproteobacteria*, Bact. = *Bacteroidetes*, Cyano. = *Cyanobacteria*, Delta. = *Deltaproteobacteria*, Firm. = *Firmicutes*, Gamma. = *Gammaproteobacteria*, Spiro. = *Spirochetes*, and ‘< 10’ = aggregated taxa that have less than 10 genomes each).

## Conclusions

Metagenomic-SIP enables us to link genome features to growth dynamics and C acquisition dynamics of bacteria as they occur in soil. We used a targeted approach, employing data from a multi-substrate DNA-SIP experiment, to select bacterial genomes that maximize life history tradeoffs. We identified genomic features (MCP, MT, OS, TF, and SMBCs) that are associated with growth and C acquisition dynamics of soil dwelling bacteria. We also identified genomic signatures (TF:gene and [SE + SM]:MT) that represent life history parameters useful in inferring bacterial ecological strategies from genome sequence data. We show that, while many intermediate strategies exist, there are diverse taxa that maximize life history tradeoffs defined by these genomic parameters. The genomic signatures we identified are readily assessed using genomic and metagenomic sequencing and these parameters may be useful in the assessment of bacterial life history strategies.

## Methods

### Soil microcosms, DNA extraction, and isopycnic centrifugation

The multi-substrate DNA-SIP experiment that provided the DNA samples we used for metagenomic-SIP has been described in detail elsewhere^15^. An overview of the experimental design for this prior DNA-SIP experiment is provided for reference in Fig. S1. Briefly, a mixture of 9 different C sources was added to soil at 0.4 mg C g^−1^ dry soil each (each representing about 3.3% of total soil C), moisture was maintained at 50% water holding capacity, and sampling was performed destructively over a period of 48 days. All treatments were derived from the exact same soil sample (from an agricultural field managed under a diverse organic cropping rotation), they received the exact same C sources, and they were incubated under the exact same conditions, the only variable manipulated was the identity of the ^13^C-labeled C source. Eight ^13^C-treatments from this prior experiment (each defined by the identity of the ^13^C source and the time of sampling) were chosen for metagenomic-SIP because the previous analysis^15^ indicated that their ^13^C-labeled DNA was enriched in bacteria that maximized differences in life history strategy (Fig. S2 and see also Fig. 5e from the prior study^15^). The treatments selected for metagenomic-SIP were: glucose day 1, xylose day 6, glucose day 14, glycerol day 14, cellulose day 30, palmitic acid day 30, palmitic acid day 48, and vanillin day 48. We also sampled ^12^C-control treatments for days 1, 6, 14, 30, and 48 to facilitate identification of ^13^C-labeled contigs and improve metagenome assembly and binning^79^. DNA used in this experiment (after undergoing extraction, isopycnic centrifugation, and fractionation) was the same as described previously^15^ and was archived at −20°C for ~2 years prior to use in this study.

### Metagenomic sequencing

For each of the eight treatments and five controls, we combined 10 μl of purified, desalted, DNA solution from each CsCl gradient fraction having a buoyant density between 1.72 and 1.77 g ml^−1^. By pooling equal volumes from these fractions, we aimed to replicate the composition of the DNA pool of the entire heavy buoyant density window (1.72-1.77 g ml^−1^). Metagenomic-SIP simulations have demonstrated that this buoyant density range sufficiently enriches for ^13^C-labeled bacterial DNA^79^. DNA amplification and sequencing were performed by the Joint Genome Institute (JGI; Berkeley, CA, USA) using standard procedures. In short, DNA was amplified and tagged with Illumina adaptors using a Nextera XT kit (Illumina Inc, San Diego, CA, USA) and sequencing was performed on the NovaSeq system (Illumina Inc).

### Read processing, metagenome assembly and annotation, and MAG binning

Quality control read processing and contig assembly was performed by the JGI as previously described^80^. Contigs were generated via terabase-scale metagenome coassembly from all 13 libraries using MetaHipMer^81^. Gene calling and annotation of assembled contigs was performed through JGI’s Integrated Microbial Genomes and Microbiomes (IMG/M) system^82^. Quality filtered reads, co-assembled contigs, and IMG annotations can be accessed through the JGI genome portal (CSP ID 503502, award DOI: 10.46936/10.25585/60000933). We mapped reads from each library to all contigs that were over 1000 bp in length using BBMap^83^ then calculated contig coverages using jgi_summarize_bam_contig_depths from MetaBAT^84^.

As we were primarily interested in genomes of bacteria that incorporated ^13^C into their DNA, we only used putatively ^13^C-labeled contigs to bin metagenome assembled genomes (MAG). Within each treatment, we defined a ^13^C-labeled contig as having an average read coverage greater than 5X in the ^13^C-treatment library and a 1.5 fold increase in coverage from the ^12^C-control to ^13^C-treatment library after accounting for difference in sequencing depths. In calculating the fold increase in coverage, we normalized for sequencing depth by dividing coverage by read counts. We binned ^13^C-labeled contigs separately for each treatment based on both tetranucleotide frequency and differential coverage with MetaBAT2^84^, MaxBin^85^, and CONCOCT^86^. Default settings were used with the exceptions that minimum contig lengths was set to 1000 bp for both MaxBin and CONCOCT and 1500 bp for MetaBAT2. Final MAGs were generated by refining bins from all three binning tools using metaWRAP^87^. Coverage information used during each binning run was from the paired ^13^C-treatment and ^12^C-control libraries, not the entire set of libraries. Therefore, we ran MAG binning eight separate times, once for each treatment. MAG qualities were calculated using CheckM^88^. For further analyses, we only used MAGs with over 50% completeness and less than 10% contamination (*i.e*., ‘medium quality’ MAGs) following the guidelines for minimum information about metagenome-assembled genomes^64^.

The binning approach we employed used co-assembled contigs, but binned these contigs separately across the eight ^13^C-labeled treatments. As such, some MAGs were identified in multiple treatments if their genomes were ^13^C-labeled by multiple ^13^C-labeled C sources. These sister MAGs might represent a single population that can derive its C from multiple C sources, or functionally distinct subpopulations each preferentially adapted for a different C source. Strain heterogeneity has previously been implicated as a cause of poor binning outcomes with soil metagenomes^89^. Traditional MAGs tend to include the entire pan-genome of heterogeneous strains representing an individual taxon^90^. Our ^13^C-labelling informed binning strategy should have greater ability to differentiate functionally differentiated sub-populations than traditional binning strategies. Further characteristics of our MAGs are discussed in Supplemental Results.

### Statistical analysis and computing

Unless otherwise stated, all statistical analyses were performed and all figures generated with R^91^ version 3.6.3. Code for all analyses and most processing is available through GitHub (https://github.com/seb369/CcycleGenomicFeatures).

### Testing associations between genomic features and activity characteristics

We first assessed associations between genomic features and activity characteristics by comparing the genetic composition of ^13^C-labeled contigs with the averaged characteristics of the ^13^C-labeled OTUs identified in each corresponding treatment from our prior study^15^. We developed a list of eight genome features hypothesized to be associated with life history strategies and microbial C-cycling activity in soil environments: 1) MCP genes were identified by the product name “methyl-accepting chemotaxis protein”. 2) Transporter genes were identified by product names containing the terms “transporter”, “channel”, “exchanger”, “symporter”, “antiporter”, “exporter”, “importer”, “ATPase”, or “pump”. The resulting gene list was then filtered to include only those predicted by TMHMM^92^ (version 2.0c) to have at least one transmembrane helix. 3) Adhesion associated genes included adhesins and holdfast and identified by product names “holdfast attachment protein HfaA”, “curli production assembly/transport component CsgG/holdfast attachment protein HfaB”, “adhesin/invasin”, “fibronectin-binding autotransporter adhesin”, “surface adhesion protein”, “autotransporter adhesin”, “adhesin HecA-like repeat protein”, “ABC-type Zn2+ transport system substrate-binding protein/surface adhesin”, “large exoprotein involved in heme utilization and adhesion”, “Tfp pilus tip-associated adhesin PilY1”, “type V secretory pathway adhesin AidA”. 4) Transcription factor genes were first identified by product names containing the terms “transcriptional regulator”, “transcriptional repressor”, “transcriptional activator”, “transcription factor”, “transcriptional regulation”, “transcription regulator”, or “transcriptional [family] regulator”, where [family] is replaced by some gene family identification. Additional transcription factor genes were identified from the protein fasta sequences using DeepTFactor^93^. 5) Osmotic stress related genes were identified by product names containing the terms “osmoregulated”, “osmoprotectant”, “osmotically-inducible”, “osmo-dependent”, “osmolarity sensor”, “ompr”, “l-ectoine synthase”. 6) Dormancy related genes covered three different mechanisms^94^. Endospore production was indicated by products containing the name “Spo0A”, though no Spo0A genes were found. Dormancy resuscitation was indicated by products containing the name “RpfC”, a resuscitation promoting factor. Dormancy related toxin-antitoxin systems were indicated by products containing the names “HipA”, “HipB”, “mRNA interferase MazF”, “antitoxin MazE”, “MazEF”, “RelB”, “RelE”, “RelBE”, “DinJ”, or “YafQ”. 7) Secreted enzyme genes were first annotated against three enzyme databases to include enzymes important for breakdown of organic matter. Carbohydrate active enzymes were annotated by mapping protein sequences to the dbCAN^95^ database (release 9.0) with HMMER using default settings. Of these enzyme genes only those in the glycoside hydrolase (GH), polysaccharide lyase (PL), or carbohydrate lyase (CE) groups were retained. Proteases were annotated by mapping protein sequences to the MEROPS^96^ database (release 12.3) using DIAMOND blastp alignment with default settings except an E-value < 1×10^−10^. Enzymes containing an α/β hydrolysis unit were annotated by mapping protein sequences to the ESTHER^97^ database (downloaded June 11^th^, 2021) with HMMER using default settings. While some enzymes containing α/β hydrolysis units are included in the carbohydrate active enzymes, this group also includes lipases. All annotated enzyme genes from these three groups were then filtered to those containing a secretion signal peptide sequence annotated by SignalP^98^ (version 5.0b). Gram + annotations were used for any genes annotated to the *Firmicutes* or *Actinobacteria* phyla, and Gram – annotations were used for all others. 8) Bacterial secondary metabolite biosynthetic gene clusters (SMBC) were predicted using antiSMASH^99^ (version 5.1.2) with default settings.

For each genomic feature, except for SMBCs, we calculated the percentage of all protein coding genes from each ^13^C-labeled contig pool (*i.e*., ^13^C-labeled in each treatment) that were annotated as described above. For SMBCs, we divided the number of SMBCs in each ^13^C-labeled contig pool by the number of protein coding genes in that pool. We then measured Pearson’s correlation between the genomic feature abundance and each of the activity characteristics averaged across the OTUs that were also ^13^C-labeled in each treatment. Within this bulk measurement, a greater percentage of the protein coding gene pool annotated to a genomic signature can indicate that, 1) a greater proportion of the represented genomes contain those genes, 2) the represented genomes have multiple copies of those genes, or 3) there is a greater diversity of those genes within the represented genomes. To account for increased false discovery rate with multiple comparisons, we adjusted *p*-values within each activity characteristic using the Benjamini-Hochburg procedure (*n* = 7).

### Examining genomic signatures of life history strategies in MAGs

We next assessed associations between genomic features and activity characteristics by comparing the genetic composition of ^13^C-labeled MAGs with the averaged characteristics of the OTUs mapping to those MAGs. As very few 16S rRNA genes were recovered and binned, we matched MAGs to ^13^C-labeled OTUs based on taxonomy and ^13^C-labeling patterns. MAG taxonomy was assigned using GTDB-Tk^65^. MAGs were taxonomically mapped to the set of OTUs that matched at the highest corresponding taxonomic level, then this set of OTUs was filtered to include those that were ^13^C-labeled in the same treatment as the MAG. Genomic features within the contigs of each MAG were determined as described above, except that for secreted enzymes, gram positive or gram negative SignalP predictions were assigned based on MAG taxonomy. Gene and SMBC counts were adjusted as before but based on the total protein coding gene count of the MAGs. We then measured Pearson’s correlation between the genomic feature abundance within the MAGs and each of the activity characteristics averaged across the OTUs mapped to the MAGs. To account for increased false discovery rate with multiple comparisons, we adjusted *p*-values within each activity characteristic using the Benjamini-Hochburg procedure (*n* = 8).

### Examining genomic signatures of life history strategies with independent studies

We analyzed publically available soil microbiome datasets to determine whether the genomic relationships we observed in ^13^C-labeld MAGs were representative of soil dwelling bacteria. Seven datasets where chosen: RefSoil^100^, Diamond *et al*. 2019^101^, Yu *et al*. 2020^102^, Wilhelm *et al*. 2019^103^, Wilhelm *et al*. 2021^104^, Zhalnina *et al*. 2018^105^, and Li *et al*. 2019^106^. Assemblies from Diamond *et al*. 2019, Yu *et al*. 2020, and Zhalnina *et al*. 2018 were downloaded from GenBank on June 21^st^, 2021 (NCBI accessions in Supplemental dataset). Assemblies from Wilhelm *et al*. 2019 and Wilhelm *et al*. 2021 were acquired from the authors. Assemblies from Li *et al*. 2019 were downloaded from figshare (https://figshare.com/s/2a812c513ab14e6c8161). Annotation was performed identically for all assemblies to avoid biases introduced by different annotation pipelines. Protein coding genes were identified and translated using Prodigal^107^ through PROKKA^108^. Transcription factor genes, SMBCs, and genes encoding transmembrane helices were further annotated as described above. Transporter genes, transcription factor genes, MCP genes, osmotic stress response genes, and SMBCs were identified and abundances were calculated as described above. 16S rRNA genes and tRNA genes were identified from PROKKA annotations. Pearson correlations were analyzed between transporter gene abundances and transcription factor gene abundances, osmotic stress response gene abundances, and SMBC abundances and between the natural log of 16S rRNA gene counts or tRNA gene counts MCP gene abundances separately for each independent dataset. Within each dataset, *p*-values were adjusted for multiple comparisons using the Benjamini-Hochburg procedure (*n* = 4).

### Using tradeoffs to define and predict life history strategies

The C-S-R framework predicts evolutionary tradeoffs in energy allocation to resource acquisition across habitats that vary temporally (*e.g*., variation in disturbance frequency). Since deletion bias in microbial genomes produces streamlined genomes of high coding density, we can assess evolutionary investment in a particular cellular system by quantifying genomic resources devoted to the operation of that system. That is, genetic information must be replicated and repaired with each generation; hence, energy allocation to a given cellular system over evolutionary time can be assessed as the proportion of the genome devoted to that system. To identify putative life history strategies for ^13^C-labeled MAGs, we used *k*-means clustering to group MAGs based genomic investment in transcription factors and resource acquisition. Investment in transcription factors was defined as the TF gene count divided by total gene counts (TF:gene). Relative investment in resource acquisition was determined by summing SE and SM counts, removing duplicates found in both categories, then dividing by the number of MT genes ([SE + SM]/MT). *k*-means clustering was performed using *k*-centroids cluster analysis with R package flexclust^109^ after scaling and centering the two values and using a *k* = 3. Statistical significance was assessed using the Kruskal-Wallis test and the Dunn test was used to assess post-hoc comparisons.

We calculated the same tradeoffs in genomic investment (TF:gene and [SE + SM]/MT) for RefSoil genomes. Predicted clusters for RefSoil genomes were made using these two genomic signatures as inferred by the R package flexclust^109^, and using the three ^13^C-labeled MAG clusters as the training dataset. Differences in genomic investments for the eight previously discussed genomic features were then assessed across clusters using the Kruskal-Wallis test with the Dunn test used to assess post-hoc comparisons. However, in this analysis, adhesion genes were identified as genes with product names containing the terms “adhesion” or “adhesins” because the previously used product names were not found in these annotations.

## Supporting information

Supplemental Results

## Acknowledgements

This material is based on work supported by the Department of Energy Office of Science, Office of Biological & Environmental Research Genomic Science Program under Award DE-SC0010558. Metagenomic sequencing, assembly, annotation, and binning (proposal: DOI 10.46936/10.25585/60000933) was performed at the U.S. Department of Energy Joint Genome Institute (https://ror.org/04xm1d337), a DOE Office of Science User Facility, supported by the Office of Science of the U.S. Department of Energy operated under Contract No. DE-AC02-05CH11231. This research was supported in part by the JGI’s Community Science Program (CSP 503502).

