## Supplemental Results for "Genomic features predict bacterial life history strategies in soil, as identified by metagenomic stable isotope probing"

*Taxonomy of ^13^C-labeled contigs*

The phylum level breakdown of the taxonomically annotated genes from the ^13^C-labeled contigs differed to some extent from the taxonomy of the ^13^C-labeled OTUs identified in Barnett *et al*. 2021^1^. (Fig. S4). For example, while *Firmicutes* made up a large portion of the ^13^C-labeled OTUs under glucose day 1, xylose day 6, and glucose day 14, very few genes from ^13^C-labeled contigs were annotated as *Firmicutes* under these treatments. Similarly, *Chloroflexi* OTUs were ^13^C-labeled under cellulose and palmitic acid day 30 and palmitic acid and vanillin day 48, yet few genes annotated to *Chloroflexi* were identified in any ^13^C-labeled contig. There are a number of potential sources for the discrepancy. First, for this metagenomic-SIP experiment we used a simple ‘heavy window’ design while Barnett *et al*. 2021^1^ used multiple-window high-resolution DNA-SIP (MW-HR-SIP). With a heavy window design, DNA from a single pooled buoyant density range is sequenced and coverage of resulting contigs is compared between the ^13^C-treatment and ^12^C-control libraries. In MW-HR-SIP, multiple overlapping heavy buoyant density windows, made up of multiple gradient fractions, are used to compare OTU abundances between a ^13^C-treatment and ^12^C-control gradient. As discussed in Barnett *et al*. 2020^2^, heavy window metagenomic-SIP may exclude low G+C DNA (*e.g., Firmicutes*) because this DNA, even when ^13^C-labeled, may be in exceedingly low abundance within the sequenced heavy window. Heavy window designs may also be less sensitive to ^13^C-labeling for high G+C DNA (*e.g.* *Actinobacteria*) because, even when unlabeled, this DNA may be found in high abundance within the sequenced buoyant density window and therefore no increase in coverage due to isotopic labeling is measured between a ^13^C-treatment library and the ^12^C-control library^2^. We chose our sequenced buoyant density window to reduce these effects but they may still be present. MW-HR-SIP may be less biased in the range of organismal G+C it can capture^3^. There is currently no comparable method to MW-HR-SIP for metagenomic sequencing though gradient resolved SIP^4^ may improve sensitivity to G+C extremes. Second, genome size may influence the apparent relative contributions of each phylum to the gene pools. Genome size can vary widely across all phyla of bacteria^5^ and larger genomes will naturally have many genes, making them appear overrepresented when assessing only genes counts within the gene pool. Third, sequencing bias, either in the 16S rRNA gene sequencing^6^ used by Barnett *et al*. 2021^1^ or in this shotgun metagenomic sequencing (*e.g.* G+C bias)^7^ may affect read recovery from particular bacterial taxa. Fourth, while the JGI IMG pipeline uses a highly diverse and inclusive reference database for metagenome annotations^8^, some taxa may be still be under-represented in the reference, limiting taxonomic annotation of their genes.

*Recovered MAGs of ^13^C-labeled bacteria*

MAGs were binned separately for each treatment-control library pair because treatments represent distinct C sources. Given this binning design, closely related MAGs can share contigs, if these MAGs were ^13^C enriched by multiple C sources (Supplemental Dataset). Since MAGs represent populations, as opposed to individual genomes, the existence of overlapping MAGs allows us to distinguish genetically similar populations that have distinct patterns of C assimilation. We recovered 27 ‘medium quality’ MAGs from the ^13^C-labeled contigs (> 50% completeness and < 10% contamination^9^; Supplemental dataset). Our binning strategy allows MAGs binned from different treatments to share co-assembled contigs. In total, MAGs encompassed 25892 contigs of which 17309 were binned into only one MAG, 7058 were binned into two MAGs, 1523 were binned into three MAGs, and 2 were binned into four MAGs. Contigs shared across MAGs tended to be small, though 210 were over 10000 bp long. We found no MAGs that were completely identical to one another (Supplemental Dataset). There were some cases where a smaller MAG appeared to be a subset of another, with up to 97.3% of its contigs shared with the larger one. When accounting for contig length, the total shared lengths were up to 98.6%. In these cases, however, less than 90% of the larger MAG, both contig and length wise, was shared with the smaller MAG. Overall, 13 MAGs shared over 50% of their contigs and length with another MAG, while 5 MAGs shared over 90% of their contigs and 6 MAGs shared over 90% of their length with other MAGs. Despite these overlaps, we believe that these MAGs likely represent distinct populations based on differences in ^13^C assimilation and growth dynamics.
