## Supplemental Tables and Figures for "Genomic features predict bacterial life history strategies in soil, as identified by metagenomic stable isotope probing"

**Table S1.** Information on the metagenomic-SIP libraries. ^13^C-labeled contigs were defined as being over 1000 bp long, having at least 5X coverage in the ^13^C-treatment library and having at least a 1.5 fold increase in coverage between the ^12^C-control and ^13^C-treatment libraries after accounting for sequencing depth. The gene counts include all genes predicted from the ^13^C-labeled contigs in each treatment.

| **Library** | **Quality**  **controlled reads** | **^13^C-labeled**  **contigs** | **Genes from**  **^13^C-labeled contigs** |
| --- | --- | --- | --- |
| ^12^C-Control day 1 | 600,398,782 |  |  |
| ^13^C-Glucose day 1 | 818,866,782 | 70,416 | 163,576 |
| ^12^C-Control day 6 | 537,771,884 |  |  |
| ^13^C-Xylose day 6 | 750,644,734 | 45,590 | 113,708 |
| ^12^C-Control day 14 | 778,966,874 |  |  |
| ^13^C-Glucose day 14 | 662,897,402 | 120,102 | 288,427 |
| ^13^C-Glycerol day 14 | 658,015,110 | 94,047 | 229,010 |
| ^12^C-Control day 30 | 1,279,583,274 |  |  |
| ^13^C-Cellulose day 30 | 707,180,514 | 214,068 | 517,605 |
| ^13^C-Palmitic acid day 30 | 632,500,566 | 194,387 | 481,889 |
| ^12^C-Control day 48 | 1,326,945,180 |  |  |
| ^13^C-Palmitic acid day 48 | 602,121,814 | 157,913 | 405,061 |
| ^13^C-Vanillin day 48 | 548,508,008 | 127,236 | 294,641 |

**Table S2**. Summary of parameter values for inferred life history clusters. A symbol (+) was added every time a cluster was observed to have a significantly higher value than another cluster in both ^13^C-MAG comparisons and the RefSoil comparisons. Abbreviations defined in the text.

|  | **MT** | **OS** | **TF** | **MCP** | **Dormancy** | **SMBC** | **SE** | **Adhesion** | ***rrn*** |
| --- | --- | --- | --- | --- | --- | --- | --- | --- | --- |
| **Ruderal** | +++^a,b^ | +++^a,b^ | ++^a,b^ | +++^a,b^ | ++^b^ |  |  |  |  |
| **Competitor** | +^b^ | ++^a,b^ | ++^a,b^ | +^b^ |  | ++^b^ | +++^a,b^ | ++^a^ |  |
| **Scarcity tolerant** |  |  |  |  |  | +^b^ | +^b^ |  |  |
| ^a^ support from ^13^C-MAG clusters  ^b^ support from RefSoil clusters | | |  |  |  |  |  |  |  |

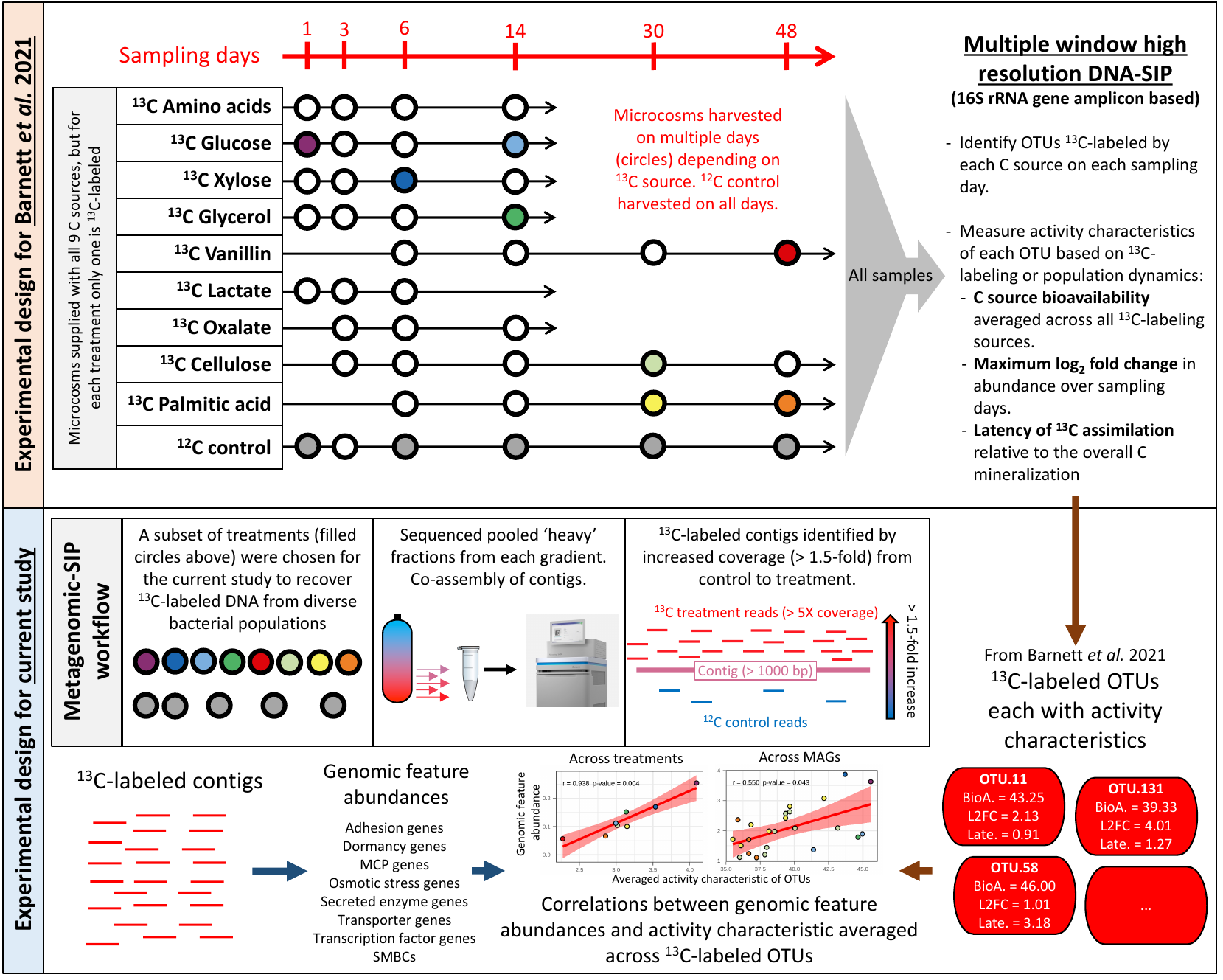

**Figure S1:** Experimental diagram of the previously described multi-substrate DNA-SIP study from Barnett *et al.,* 2021 (top panel) and the newly described metagenomic-SIP sequencing, processing, and analysis described here (bottom panel). In the top panel, the circles represent the days when microcosms were harvested, while the filled circles represent the samples chosen for metagenomic-SIP sequencing. The original DNA-SIP study used multiple-window high-resolution DNA-SIP to identify bacterial operational taxonomic units (OTU) that assimilated ^13^C from each of the ^13^C sources within the harvested microcosms. The ^13^C-labeling patterns along with population dynamics in soils (unfractionated DNA) were used to generate the three activity characteristics. To compare genomic features in ^13^C-labeled contigs within a treatment or MAG, we then averaged the characteristics of the OTUs either ^13^C-labeled in the treatment or taxonomically mapped to the MAG and ^13^C-labeled in the same treatment respectively. For the metagenomic-SIP sequencing, pooled fractions between 1.72 and 1.77 g ml^-1^ were sequenced. ^13^C-labeled contigs were distinguished as being > 1000 bp long, having at least 5X coverage in the ^13^C-treatment library, and having over a 1.5 fold increase in coverage in the ^13^C-treatment compared to its corresponding ^12^C-control.

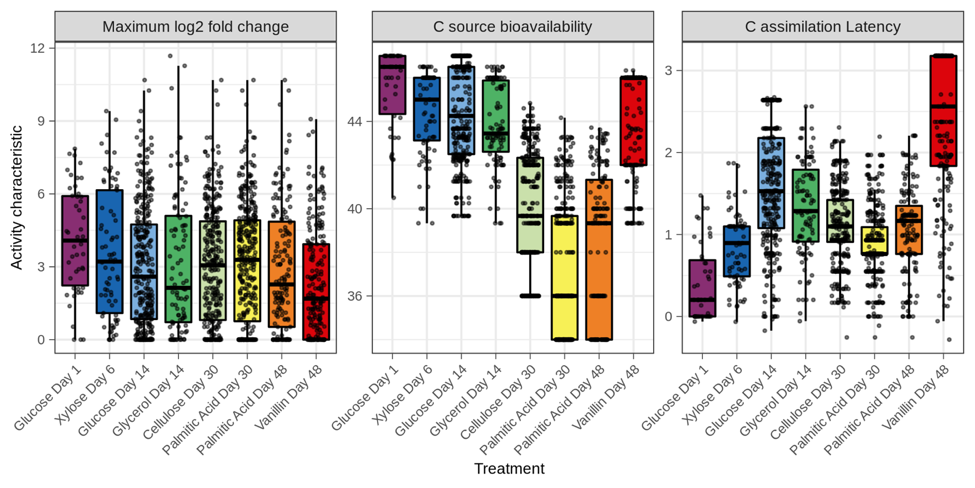

**Figure S2:** Activity characteristic values of the ^13^C-labeled OTUs (as previously described in Barnett *et al.* 2021) detected in each ^13^C-labeled treatment subjected to metagenomic sequencing. Briefly, maximum log_2_ fold change represents the change in differential abundance from the initial condition (time zero) to the point when relative abundance was maximal for a given OTU. C source bioavailability is defined as the average bioavailabilty of all the substrates assimilated by each OTU, with the bioavailability of each C source defined operationally based on its mineralization dynamics (as previously described in Barnett *et al.,* 2021). C assimilation latency is defined as the difference in time between the point when ^13^C source mineralization was maximal and the point at which an OTU was observed to assimilate the ^13^C-substrate.

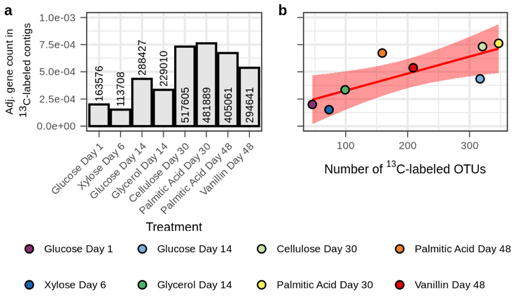

**Figure S3:** The number of genes in ^13^C-labeled contigs correlates with the number of ^13^C-labeled OTUs identified in Barnett *et al.,* 2021, suggesting that we recovered genomic content of the target, active bacteria. Gene count is normalized by the number of reads recovered from the ^13^C-treatment libraries (Table S1). **a)** The normalized number of genes found in the ^13^C-labeled contigs from each treatment. The numbers within or above the bars indicate the number of genes before normalization. **b)** The relationship between the normalized number of genes in the ^13^C-labeled contigs and the number of OTUs ^13^C-labeled under the same treatment (Pearson’s *r* = 0.795, *p-*value = 0.018). The red line represents the linear regression with red shading indicating the 95% confidence intervals.

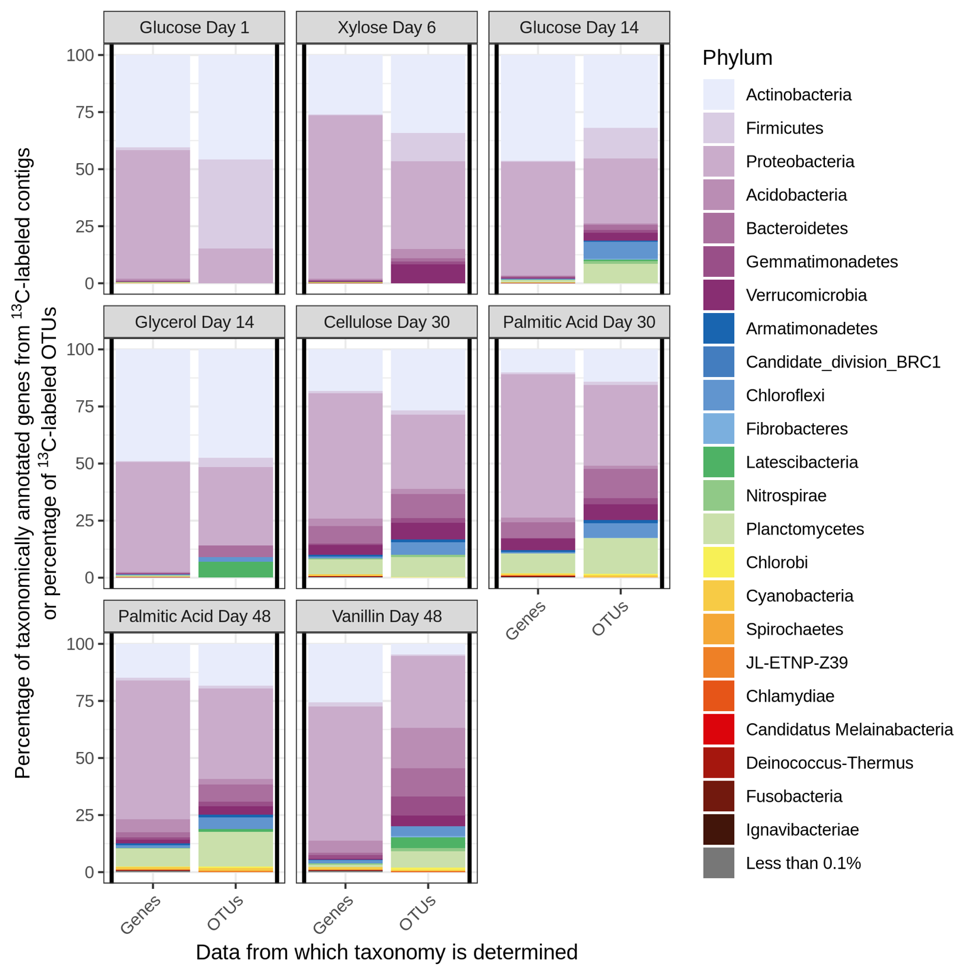

**Figure S4:** Phylum level breakdown of the taxonomically annotated genes in the ^13^C-labeled contigs under each treatment (Genes) and the phylum level breakdown of the ^13^C-labeled OTUs under each treatment in Barnett *et al.*, 2021 (OTUs). Genes used for this analysis were only those with taxonomic annotations. Gene taxonomy was assigned using the IMG pipeline. OTU taxonomy was assigned using the SILVA 111 release.

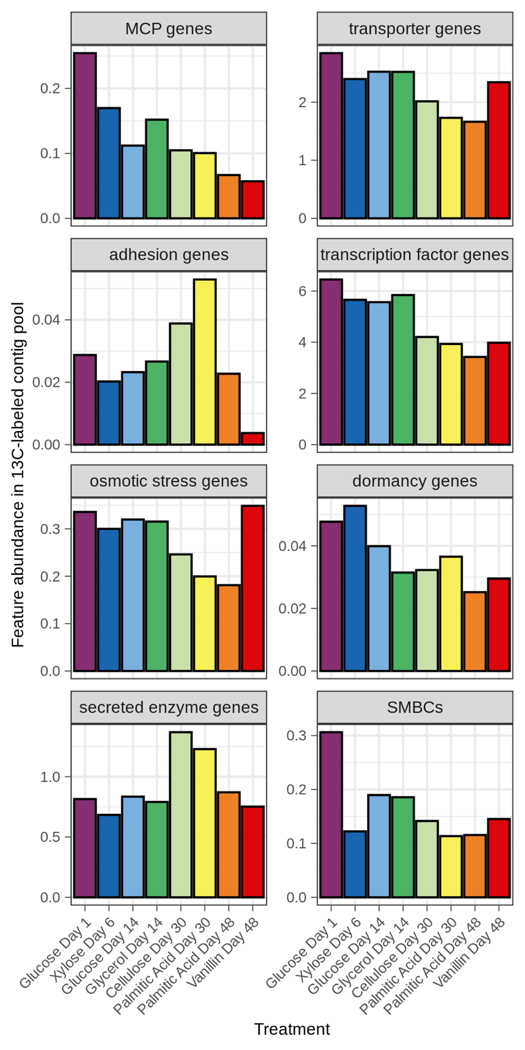

**Figure S5:** Abundance of genes from each feature in the ^13^C-labeled contigs from each treatment. For all features except SMBCs, feature abundance is calculated as the percentage of total protein coding genes having the feature. For SMBCs, abundance is calculated as the number of SMBCs divided by the total protein coding gene count.

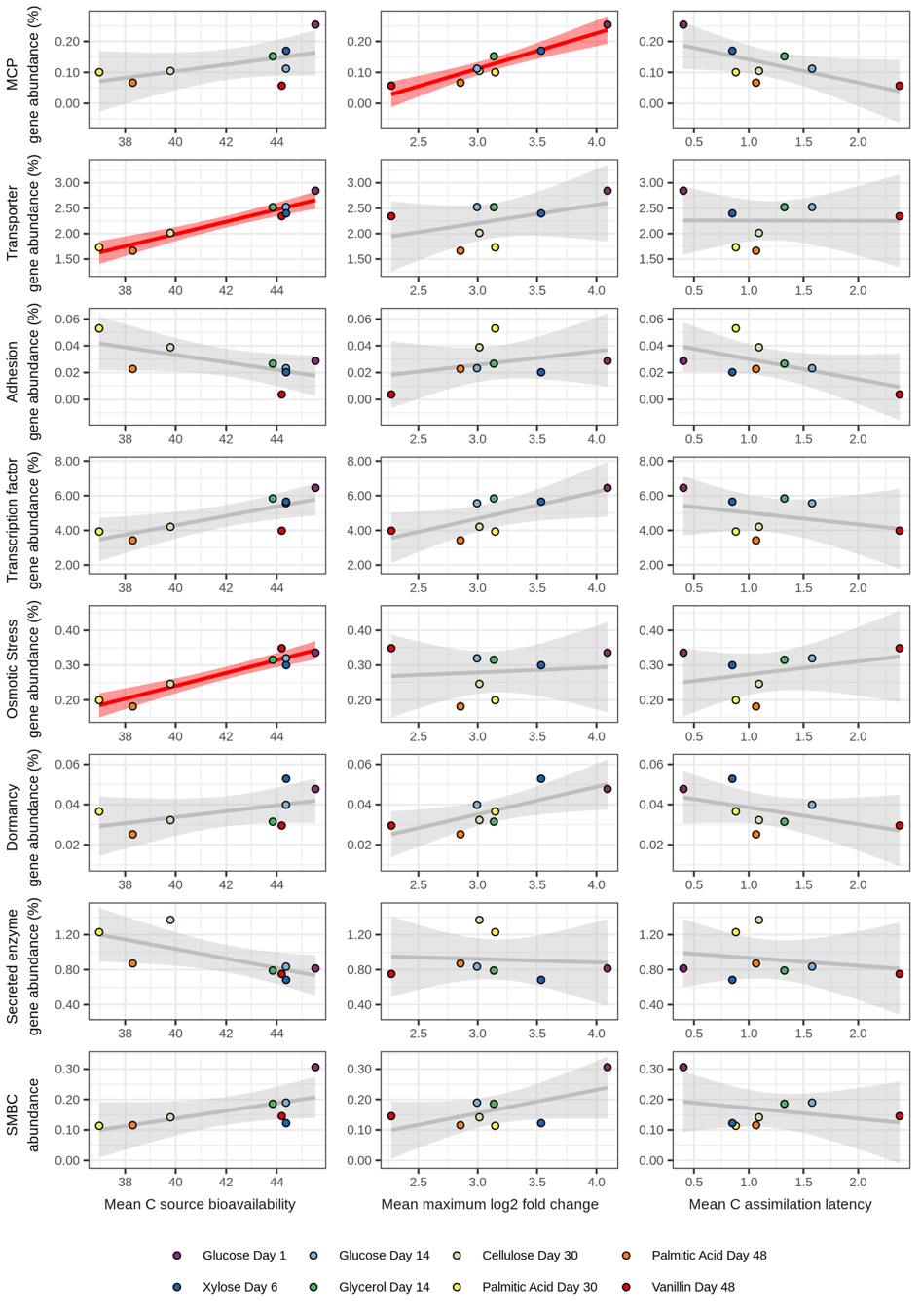

**Figure S6:** Relationships between the frequency of 8 genome features in ^13^C-labeled contigs and all three *in situ* activity characteristics of ^13^C-labeled OTUs across treatments. For all except SMBCs, abundance is calculated as the percent of protein coding genes in ^13^C-labeled contigs that are annotated within the genomic feature. SMBC abundance is calculated as the SMBC count divided by total protein coding gene count. Red or grey lines represent the linear relationships with shading indicating the 95% confidence intervals. Red relationships are statistically significant, with p-values adjusted for multiple comparisons using the Benjamini-Hochburg procedure (*n* = 8). Correlation statistics are listed in the Supplemental Dataset.

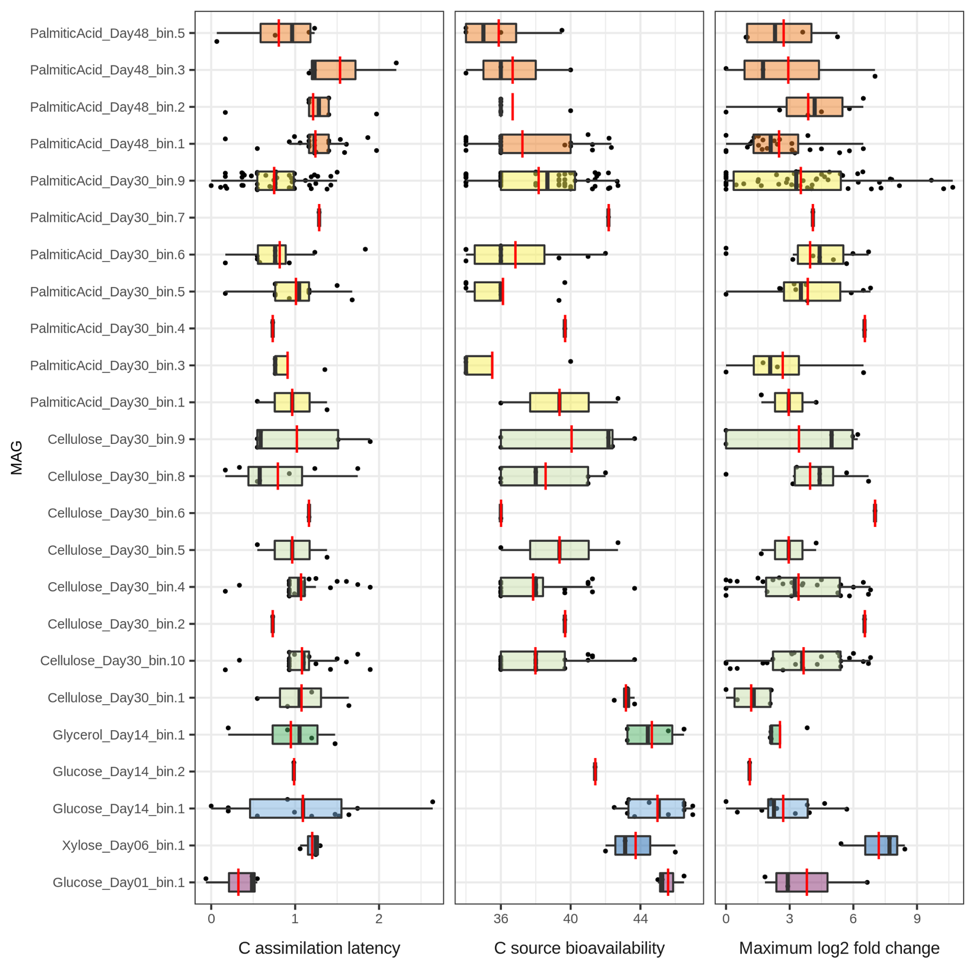

**Figure S7:** Activity characteristics of the ^13^C-labeled OTUs mapped to each ^13^C-labeled MAG. Boxplots are colored by the treatment under which ^13^C-labeling occurs. Red bars indicate mean values. Three MAGs had no matching OTUs: Cellulose_Day30_bin.7, PalmiticAcid_Day48_bin.4, and Vanillin_Day48_bin.1. MAG and mapped OTU details are found in the Supplemental Dataset.

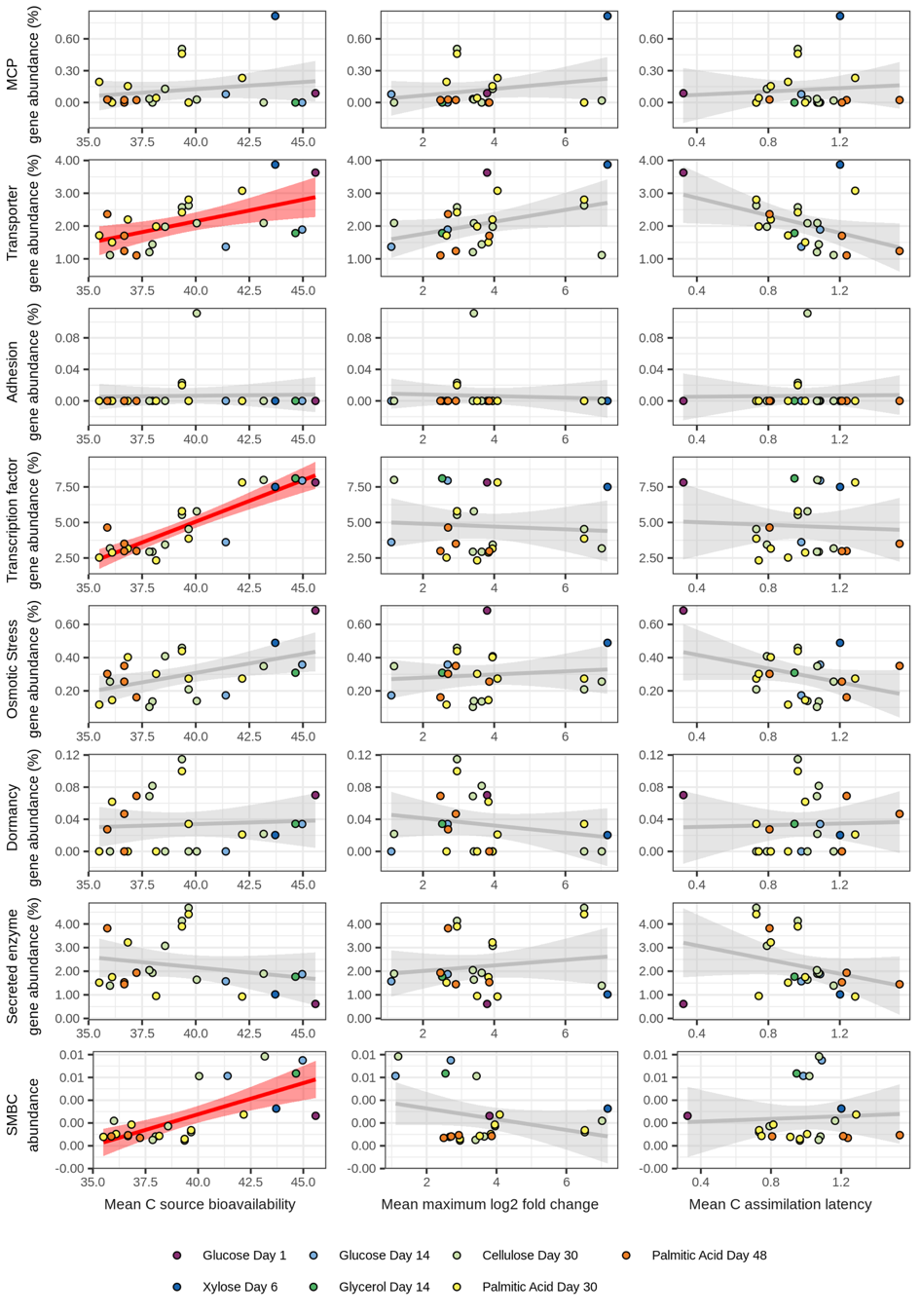

**Figure S8:** Relationships between abundance of all 8 genome features in ^13^C-labeled MAGs and all three activity characteristics of OTUs matching MAG taxonomy and ^13^C-labeling. For all except SMBCs, abundance is calculated as the percent of protein coding genes in the MAG that are annotated within the genomic feature. SMBC abundance is calculated as the SMBC count divided by total protein coding gene count. Red or grey lines represent the linear relationships with shading indicating the 95% confidence intervals. Red relationships are statistically significant, with *p*-values adjusted for multiple comparisons using the Benjamini-Hochburg procedure (*n* = 8). Correlation statistics are listed in the Supplemental Dataset.

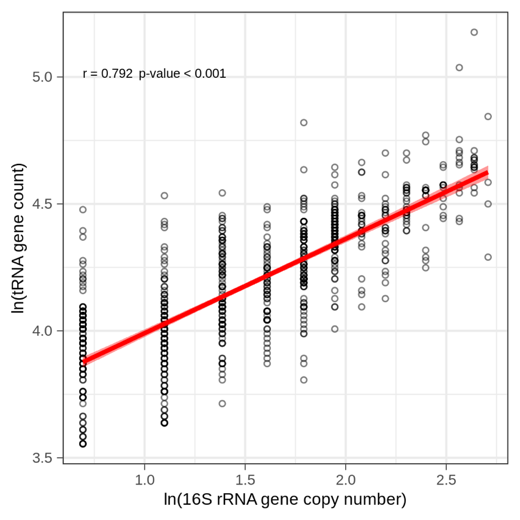

**Figure S9:** Relationship between the natural log of 16S rRNA gene counts and tRNA gene counts across the RefSoil genomes. Pearson’s *r* and *p*-value are displayed in plot. The red line represents the linear relationships with shading indicating the 95% confidence interval.

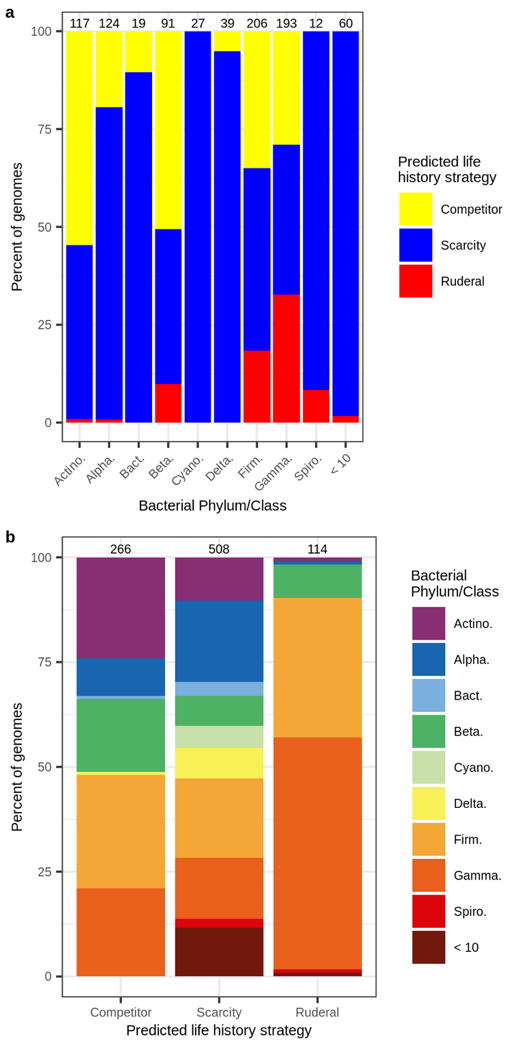

**Figure S10:** Distribution of RefSoil bacterial taxa (at the phylum or class level) across the life history strategies predicted from clusters of TF:gene and [SE + SM]:MT. **a)** Percentage of genomes from each taxa in each predicted life history strategy with total number of genomes used above each stacked bar. **b)** Percentage of genomes in each predicted life history strategy cluster that are classified to each phylum/class. Phylum/class abriviations are Actino. = *Actinobacteria*, Alpha. = *Alphaproteobacteria*, Bact. = *Bacteroidetes*, Cyano. = *Cyanobacteria*, Delta. = *Deltaproteobacteria*, Firm. = *Firmicutes*, Gamma. = *Gammaproteobacteria*, Spiro. = *Spirochetes*, and ‘< 10’ = taxa that contain less than 10 genomes.

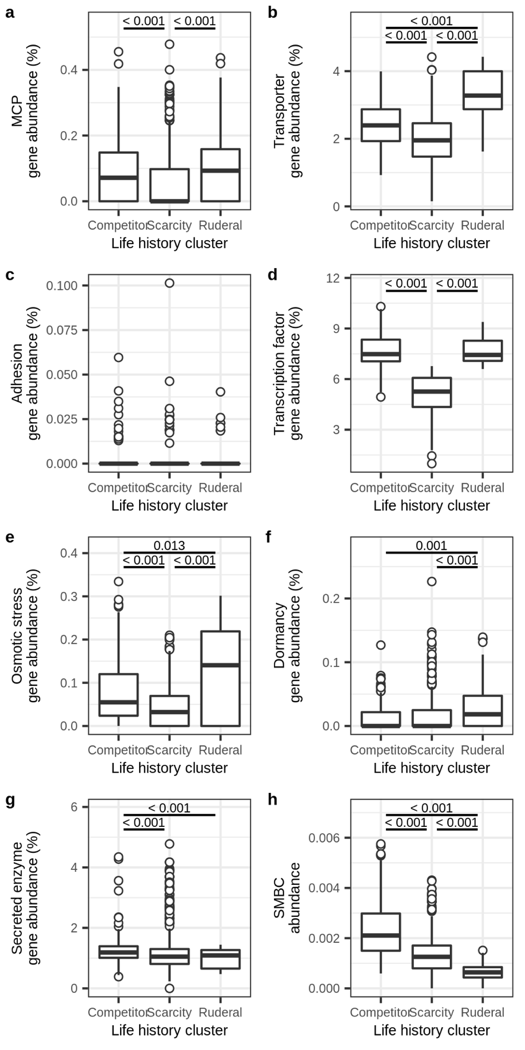

**Figure S11:** Genomic investment in gene systems differ across life history strategies predicted from TF:genes and [SE + SM]:MT. Data are from RefSoil genomes with *k*-means clustering trained by clusters identified from ^13^C-labeled MAG. In all cases variation across life history clusters was first tested with Kruskal-Wallis tests and where statistically significant (*p*-value < 0.05) post hoc pairwise tests were performed using Dunn tests. **a)** Scarcity adapted genomes have a lower investment in MCP genes than competitor or ruderal adapted genomes. **b)** Ruderal genomes have a higher investment in MT than competitor or scarcity adapted genomes and competitor adapted genomes have higher investment than scarcity adapted genomes. **c)** There is no statistically significant difference in investment in adhesion genes across clusters. **d)** Scarcity adapted genomes have lower investment in TF than ruderal or competitor adapted genomes. **e)** Ruderal genomes have a higher investment in osmotic stress response genes than competitor or scarcity adapted genomes and competitor adapted genomes have higher investment than scarcity adapted genomes. **f)** Ruderal genomes have a higher investment in dormancy genes than competitor or scarcity adapted genomes **g)** Competitor genomes have a higher investment in SE than ruderal or scarcity adapted genomes. **h)** Competitor genomes have a higher investment in SMBCs than ruderal or scarcity adapted genomes and scarcity adapted genomes have higher investment than ruderal genomes.
